# Lunaemycins, new cyclic hexapeptide antibiotics from the cave moonmilk-dweller *Streptomyces lunaelactis* MM109^T^

**DOI:** 10.1101/2022.12.15.520542

**Authors:** Loïc Martinet, Aymeric Naômé, Lucas C. D. Rezende, Déborah Tellatin, Bernard Pignon, Jean-Denis Docquier, Filomena Sannio, Dominique Baiwir, Gabriel Mazzucchelli, Michel Frédérich, Sébastien Rigali

## Abstract

*Streptomyces lunaelactis* strains have been isolated from moonmilk deposits which are calcium carbonate speleothems used for centuries in traditional medicine for their antimicrobial properties. Genome mining revealed that these strains are a remarkable example of a *Streptomyces* species with huge heterogeneity regarding their content in biosynthetic gene clusters (BGCs) for specialized metabolite production. BGC 28a is one of the cryptic BGCs that is only carried by a subgroup of *S. lunaelactis* strains for which *in silico* analysis predicted the production of nonribosomal peptide antibiotics containing the non-proteogenic amino acid piperazic acid (Piz). Comparative metabolomics of culture extracts of *S. lunaelactis* strains either or not holding BGC 28a combined with MS/MS-guided peptidogenomics and ^1^H/^13^C NMR allowed to identify the cyclic hexapeptide with the amino acid sequence (D-Phe)-(L-HO-Ile)-(D-Piz)-(L-Piz)-(D-Piz)-(L-Piz), called lunaemycin A, as the main compound synthesized by BGC 28a. Molecular networking further identified 18 additional lunaemycins, 14 of them having their structure elucidated by HRMS/MS. Antimicrobial assays demonstrated a huge bactericidal activity of lunaemycins against Gram-positive bacteria including multi-drug resistant clinical isolates. Our work demonstrates how accurate *in silico* analysis of a cryptic BGC can highly facilitate the identification, the structural elucidation, and the bioactivity of its associated specialized metabolites.

## 1. Introduction

*Streptomyces* and other actinomycetes are Gram-positive filamentous bacteria that offered most of the molecules of microbial origin which are now used in human and animal therapy [1,2]. The urge for new structural leads for both the agro-food and pharmaceutical fields has revitalized the interest for bioprospection of microorganisms dwelling in the most diverse and extreme environmental niches [3]. This quest for metabolite-producing bacteria in environments far from the ecological niches where they were mainly/originally isolated (such as rich organic soils for actinomycetes) is motivated by the idea that they would possess genetic features adapted to these specific and unusual environments allowing them to produce uncommon ‘specialized’ metabolites [4].

Caves, being inorganic and extreme oligotrophic environments, were initially considered as inhospitable places for streptomycetes and other microorganisms programmed for primarily consuming nutrients derived from the plant decomposing organic matter. Instead, this phylum is surprisingly omnipresent, displaying relative abundance and diversity depending on the type of cave and speleothem [5–10]. One of such geological formations previously unexpected to house a huge diversity of actinomycetes is the so-called moonmilk, a soft white carbonate deposit in limestone caves mainly consisting of microfibers of calcite [11–13]. Even within the same cave, the diversity of actinomycetes can widely differ from one moonmilk deposit to another [6]. More astonishingly, these bacteria do not only inhabit moonmilk to benefit from the nutrients carried by the water percolating from the surface, but they also actively participate in the creation of the mineral structure itself [12,14–18].

The decision of our laboratory to search for antibiotic-producing bacteria in moonmilk deposits originated from evidence of hundreds of years of mining activities for the use of this speleothem as powder with antimicrobial properties [13,19]. The first scanning electron micrographs of the calcite fibers of moonmilk also revealed filamentous bacteria further supporting the presence of microorganisms notorious as ‘antibiotic-makers’ [14–16,18]. And indeed, each actinomycete we isolated from three different moonmilk deposits of the cave ‘la grotte des collemboles’ (Comblain-au-Pont, Belgium) displayed antibacterial and/or antifungal activities [9,10]. A *Streptomyces* species that we found in all moonmilk deposits of the cave is *Streptomyces lunaelactis* with isolate MM109^T^ as the archetype strain [9,10,20,21]. *S. lunaelactis* MM109^T^ is the first fully characterized and completely sequenced *Streptomyces* strain originating from these speleothems [20–22]. Genome mining of the 18 *S. lunaelactis* strains revealed 42 biosynthetic gene clusters (BGC) with only 18 of them being conserved among all strains thereby providing a remarkable example of a huge specialized metabolism heterogeneity within a single species [22].

One of our aims in investigating cave microbiology is to identify the genetic and physiological features that make soil-dwelling bacteria to adapt to inorganic mineral environments [15]. Some of these adjustments must be part of their specialized metabolism used to capture essential nutrients and compete with other microorganisms or instead contribute to the survival of the entire microbial community. In our effort at identifying the metabolites produced by moonmilk-dwelling actinomycetes, interesting findings came to light such as the discovery of the BGC responsible for the hallmark green pigmentation of *S. lunaelactis* strains which revealed a unique example of a BGC involved in the production of two structurally and functionally different metabolites, namely the bagremycin antibiotics and the iron chelating molecules that generate ferroverdins [22–24].

Herein, we report how genome mining guided MS/MS-based networking and peptidogenomics allowed the discovery, the elucidation of the structure and the biosynthetic pathway of lunaemycins, new potent antibiotics produced by *S. lunaelactis* MM109^T^. Lunaemycins are new piperazic acid-containing cyclic hexapeptides able to inhibit the growth of all tested Gram-positive bacteria including various multi-drug resistant isolates, thereby in part explaining the success of this species at colonizing extremely competitive environments.

## 2. Results and Discussion

### 2.1. Sequence analysis of BGC 28a and predicted structure of its associated natural product

Mining the genome of *S. lunaelactis* MM109^T^ [21] with antiSMASH identified BGC 28a, a ~69 kb non-ribosomal peptide synthetase (NRPS)-type BGC consisting of 39 open reading frames (ORFs) from nucleotide positions 3,124 to 72,040 on the linear plasmid pSLUN1a (**Figure 1** and **Table 1**). The *hmt* gene cluster of *Streptomyces himastatinicus* producing the antitumor antibiotic himastatins [25,26] is the most similar BGC of the MIBiG database [27] in terms of number of homologous genes (**Figure S1** and **Table 1**). Further sequence analysis highlighted three other known BGCs sharing common features with BGC 28a, namely the *ktz* gene cluster producing the antifungal kutznerides in *Kutzneria* sp. 744 [28], the *sten* gene cluster associated with the production of the antimicrobial stenothricins in *Streptomyces roseosporus* NRRL 15998 [29], and the *art* BGC for the antitumor antibiotic aurantimycins produced by *Streptomyces aurantiacus* JA 4570 [30]. The synteny and conserved gene content of these four BGCs is relatively low with *hmt*, *ktz*, *art*, and *sten* gene clusters sharing only nine, seven, six, and four homologous genes with the 39 ORFs of BGC 28a, respectively (**Figure S1**). Himastatins, kutznerides, stenothricins, and aurantimycins are all cyclic depsipeptides synthesized by NRPS or hybrid NRPS-PKS machineries [31–34], suggesting that the main product of BGC 28a should partially share some structural features and/or amino acid building blocks with these natural products.

**Figure 1.**
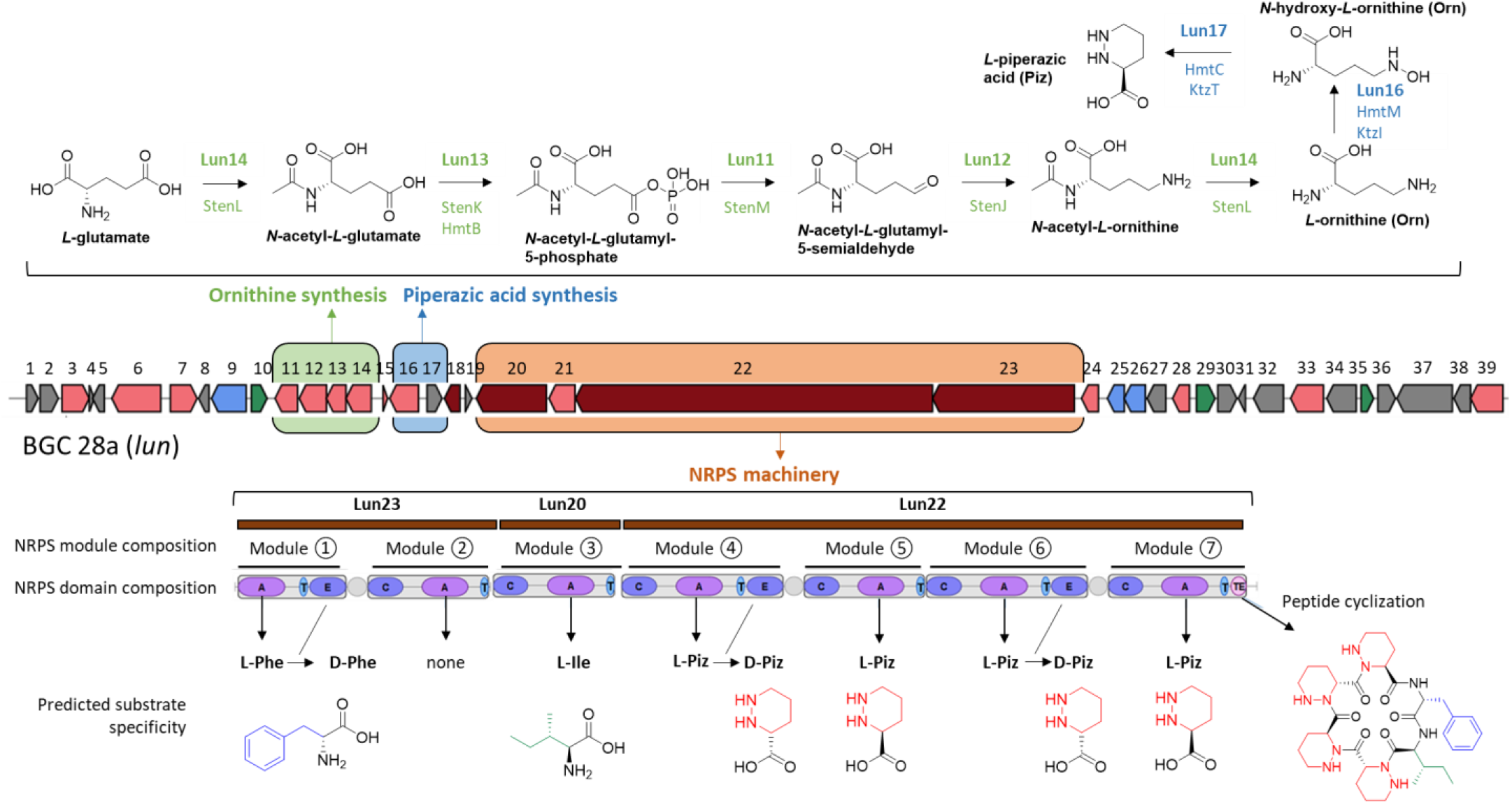
Predicted biosynthetic pathways and substrate specificity of BGC 28a (*lun* cluster). Top: genes/proteins predicted to be involved in the Ornithine (Orn) and Piperazic acid (Piz) biosynthesis. Bottom: modules, domains, predicted amino acid building-blocks, and modifying reactions of NRPSs of BGC 28a predicted to generate a cyclized hexapeptide with a monoisotopic mass of 708.4041 Da. Abbreviations: A, adenylation domain; C, condensation domain; T, thiolation domain; TE, thioesterase domain; Ile, isoleucine; Phe, phenylalanine; Piz, piperazic acid.

**Table 1.** Deduced functions of ORFs of the *lun* (BGC 28a) biosynthetic gene cluster of *S. lunaelactis* MM109^T^.

| ORFs |  | Closest homologue in MiBIG database (Protein ID) | Homology (id/cov) | Proposed function |
| --- | --- | --- | --- | --- |
| Locus Tag | Name |  |  |  |
| Accession no. | size (aa) |  |  |  |
| SLUN_RS38410<br>WP_108155188.1 | Lun1<br>177 | TmcL, hypothetical protein, $\alpha,\beta$-epoxyketone BGC of *Streptomyces chromofuscus* (CUX96959.1) | 64/100 | Unknown |
| SLUN_RS38415<br>WP_108155189.1 | Lun2<br>278 | DUF4097 family beta strand repeat-containing protein, $\alpha,\beta$-epoxyketone BGC of *Streptomyces chromofuscus* (CUX96960.1) | 48/100 | Unknown |
| SLUN_RS38420<br>WP_108155190.1 | Lun3<br>416 | Cytochrome P450, streptovaricin BGC of *Streptomyces spectabilis* (ASZ00144.1) | 61/97 | product modification |
| SLUN_RS38425<br>WP_108155191.1 | Lun4<br>76 | Ferredoxin, caniferolide A BGC from *Streptomyces caniferus* (QBF51761.1) | 52/82 | product modification |
| SLUN_RS38430<br>WP_254708284.1 | Lun5<br>179 | GtrA family protein, monensin BGC from *Streptomyces cinnamomensis* (AAO65788.1) | 60/82 | Unknown |
| SLUN_RS38435<br>WP_159100457.1 | Lun6<br>746 | MPPL family transporter, alnumycin A BGC from *Streptomyces* sp. CM020 (ACI88887.1) | 58/98 | Exporter |
| SLUN_RS38440<br>WP_108155192.1 | Lun7<br>409 | MfnN, cytochrome P450, marformycin A BGC from *Streptomyces drozdowiczii* (AJV88386.1) | 37/100 | product modification |
| SLUN_RS38445<br>WP_108155193.1 | Lun8<br>157 | Hypothetical protein, no hit in MiBIG, closest to *Streptomyces* sp. Ag82_O1-15 (WP_095849664.1) | 47/88 | Unknown |
| SLUN_RS38450<br>WP_108155194.1 | Lun9<br>531 | MFS transporter, siamycin I BGC from *Streptomyces* sp. (BBD82029.1) | 72/93 | Resistance export |
| SLUN_RS38455<br>WP_108155195.1 | Lun10<br>241 | TetR/AcrR family transcriptional regulator, siamycin I BGC from *Streptomyces* sp. (BBD82030.1) | 61/96 | Resistance regulation |
| SLUN_RS38460<br>WP_108155196.1 | Lun11<br>342 | StenM, N-acetyl-gamma-glutamyl-phosphate reductase (ArgC), stenothricin BGC from *Streptomyces roseosporus* (EFE73305.1) | 81/100 | L-ornithine synthesis |
| SLUN_RS38465<br>WP_108155197.1 | Lun12<br>417 | StenJ, N2-acetyl-L-ornithine:2-oxoglutarate aminotransferase (ArgD), stenothricin BGC from *Streptomyces roseosporus* NRRL 15998 (EFE73302.1) | 72/94 | L-ornithine synthesis |
| SLUN_RS38470<br>WP_108155198.1 | Lun13<br>300 | StenK, N-acetylglutamate kinase (ArgB), stenothricin BGC from *Streptomyces roseosporus* (EFE73303.1) | 81/97 | L-ornithine synthesis |
| SLUN_RS38475<br>WP_108155199.1 | Lun14<br>384 | StenL, bifunctional N2-acetyl-L-ornithine:L-glutamate N-acetyltransferase (ArgJ), stenothricin BGC from *Streptomyces roseosporus* (EFE73304.1) | 80/100 | L-ornithine synthesis |
| SLUN_RS38480<br>WP_108155200.1 | Lun15<br>70 | KtzJ, aurantimycin A BGC from *Streptomyces aurantiacus* JA 4570 (WP_016638464.1) | 71/100 | Building block modification |
| SLUN_RS38485<br>WP_108155201.1 | Lun16<br>444 | HmtM, lysine N(6)-hydroxylase/L-ornithine N(5)-oxygenase, himastatin BGC of *Streptomyces himastatinicus* (CBZ42147.1) | 63/97 | Piz synthesis |
| SLUN_RS38490<br>WP_108155202.1 | Lun17<br>230 | HmtC, Piperazate synthase, himastatin BGC *Streptomyces himastatinicus* (CBZ42137.1) | 69/95 | Piz synthesis |
| SLUN_RS38495<br>WP_108155203.1 | Lun18<br>235 | LmbU family transcriptional regulator, himastatin BGC *Streptomyces himastatinicus* (CBZ42138.1) | 69/95 | Regulation |
| SLUN_RS38500<br>WP_108155204.1 | Lun19<br>100 | chorismate mutase, no hit in MiBIG, *Streptomyces* sp. SLBN-118 (WP_142214144.1) | 78/97 | Phe biosynthetic pathway |
| SLUN_RS38505<br>WP_108155205.1 | Lun20<br>1072 | OciC, Non-ribosomal peptide synthetase (Module 3, domain C-A-T), cyanopeptolin BGC from *Planktothrix agardhii* NIVA-CYA 116 (ABI26079.1) | 43/98 | Incorporated aa:<br>L-Ile |
| SLUN_RS38510<br>WP_108155206.1 | Lun21<br>400 | NAD(P)/FAD-dependent oxidoreductase, aurantimycin A BGC from *Streptomyces aurantiacus* JA 4570 (WP_016638468.1) | 53/99 | hydroxylation |
| SLUN_RS38515<br>WP_108155207.1 | Lun22<br>5462 | HmtL, Non-ribosomal peptide synthetase (Modules 4/5/6/7, domains C-A-T-E/C-A-T/C-A-T-E/C-A-T-TE), himastatin BGC of *Streptomyces himastatinicus* (CBZ42146.1) | 55-100 | Incorporated aa:<br>D-Piz/L-Piz/D-Piz/L-Piz and peptide release and cyclization |
| SLUN_RS38520<br>WP_159100458.1 | Lun23<br>2167 | Non-ribosomal peptide synthetase (Module 1/2, domains A-T-E/C-A-T), aurantimycin A BGC from *Streptomyces aurantiacus* JA 4570 (WP_016638470.1) | 53/100 | Incorporated aa:<br>D-Phe/inactive |
| SLUN_RS38525<br>WP_108155209.1 | Lun24<br>259 | KtzF thioesterase, alpha/beta fold hydrolase, himastatin BGC of *Streptomyces himastatinicus* (CBZ42144.1) | 58/97 | Peptide release |
| SLUN_RS38530<br>WP_108155210.1 | Lun25<br>258 | ABC transporter permease, aurantimycin A BGC from *Streptomyces aurantiacus* JA 4570 (WP_016638474.1) | 51/100 | Drug resistance |
| SLUN_RS38535<br>WP_108155211.1 | Lun26<br>323 | Daunorubicin resistance protein DrrA family ABC transporter ATP-binding protein, aurantimycin A BGC of *Streptomyces aurantiacus* JA 4570 (WP_016638473.1) | 66/94 | Drug resistance |
| SLUN_RS38540<br>WP_108155212.1 | Lun27<br>286 | DUF4097 family beta strand repeat-containing protein, himastatin BGC *Streptomyces himastatinicus* (CBZ42150.1) | 39/100 | Unknown |
| SLUN_RS38545<br>WP_108155213.1 | Lun28<br>261 | 3-oxoacyl-ACP reductase FabG, kiamycin biosynthetic gene cluster from *Streptomyces* sp. W007 (EHM27508.1) | 74/97 | Unknown |
| SLUN_RS38550<br>WP_108155214.1 | Lun29<br>285 | SARP family transcriptional regulator, mitomycin biosynthetic gene cluster from *Streptomyces lavendulae* (AAD28451.1) | 30/90 | Regulation |
| SLUN_RS38555<br>WP_108155215.1 | Lun30<br>293 | class I SAM-dependent methyltransferase, 5-isoprenylindole-3-carboxylate β-D-glycosyl ester BGC from *Streptomyces* sp. RM-5-8 (ANA09433.1) | 31/85 | Unknown |
| SLUN_RS38560<br>WP_108155216.1 | Lun31<br>108 | ArsR family transcriptional regulator, no hit in MiBIG database, *Streptomyces mangrovisoli* (WP_052743136.1) | 93/85 | Regulation |
| SLUN_RS38565<br>WP_108155217.1 | Lun32<br>457 | M14 family zinc carboxypeptidase, capreomycin IA BGC from *Saccharothrix mutabilis* subsp. Capreolus (ABR67765.1) | 45/98 | Unknown |
| SLUN_RS38570<br>WP_108155218.1 | Lun33<br>500 | Amino acid permease, amphotericin B biosynthetic gene cluster from *Streptomyces nodosus* (AAV48836.1) | 54/89 | Unknown |
| SLUN_RS38575<br>WP_257153969.1 | Lun34<br>450 | glutamine synthetase family protein, colabomycin E BGC from *Streptomyces aureus* (AIL50155.1) | 36/100 | Unknown |
| SLUN_RS38580<br>WP_108155220.1 | Lun35<br>190 | MarR family transcriptional regulator, no hit in MiBIG database, *Streptomyces* sp. ISL-44 (WP_215144797.1) | 83/93 | Regulation |
| SLUN_RS38585<br>WP_108155221.1 | Lun36<br>268 | acetoacetate decarboxylase family protein, enduracidin BGC from *Streptomyces fungicidicus* (ABD65946.1) | 35/99 | Unknown |
| SLUN_RS38590<br>WP_108155222.1 | Lun37<br>846 | SpoIIE family protein phosphatase, laidlomycin BGC from *Streptomyces* sp. CS684 (AFL48546.1) | 48/68 | Unknown |
| SLUN_RS38595<br>WP_108155223.1 | Lun38<br>256 | gamma-glutamyl-gamma-aminobutyrate hydrolase, nenestatin biosynthetic gene cluster from *Micromonospora echinospora* (ARD70860.1) | 35/95 | Unknown |
| SLUN_RS41955<br>WP_108155224.1 | Lun39<br>495 | Putative aldehyde dehydrogenase, pederin BGC from uncultured bacterium (AAS47555.1) | 49/100 | Unknown |

Closer sequence analysis of core and additional biosynthetic genes of BGC 28a allowed to propose a biosynthetic pathway and predict the amino-acid building blocks and backbone of its main product (**Figure 1**). First, as observed in the BGCs of kutznerides and himastatins, BGC 28a contains the two genes required for the biosynthesis of the non-proteinogenic amino acid piperazic acid (Piz) from L-ornithine (Orn). The first reaction is performed by the product of *lun16* encoding the flavin- and oxygen-dependent L-ornithine N-hydroxylase (SLUN_ RS38485 homologous to HmtM and KtzI, **Figure 1**, **Table 1**) to give L-N^*5*^-OH-ornithine [35], that will be further converted to L-Piz by the product of *lun17* (SLUN_ RS38490 homologous to HmtC and KtzT, **Figure 1**, **Table 1**) encoding the heme-dependent piperazate synthase [36]. Interestingly, genes encoding piperazate synthase are present in most BGCs for Piz-containing molecules [37,38], suggesting that at least one NRPS module of BGC 28a is expected to recruit Piz as building block. Next, as observed in the stenothricin gene cluster [29], BGC 28a contains a transcription unit with four ORFs whose products encode the enzymes involved in the five steps of Orn biosynthesis from glutamic acid and glutamine precursors (**Figure 1**, **Table 1**). These genes/proteins are *lun14* for the bifunctional N^2^-acetyl-L-ornithine:L-glutamate N-acetyltransferase (SLUN_ RS38475 homologous to StenL, **Figure 1**, **Table 1**), *lun13* for the N-acetylglutamate kinase (SLUN_RS38470 homologous to StenK and HmtB, **Figure 1**, **Table 1**), *lun11* for the N-acetyl-gamma-glutamyl-phosphate reductase (SLUN_RS38460 homologous to StenM, **Figure 1**, **Table 1**), and *lun12* for the N^2^-acetyl-L-ornithine:2-oxoglutarate aminotransferase (SLUN_RS38465 homologous to StenJ, **Figure 1**, **Table 1**). BGC 28a thus contains all genes required for the enzymatic conversion of glutamic acid and glutamine to L-Piz, further supporting the recruitment of this amino acid by the NRPS machinery.

Regarding the core biosynthetic genes, three ORFs encode NRPSs with complete biosynthetic modules. The sequential order of these modules in peptide synthesis (initiation, elongation, and termination modules) is predicted based on the composition of their catalytic domains (**Figure 1**). The biosynthesis would start with Lun23 (SLUN_ RS38520, providing the initiation module 1, and the ‘inactive’ module 2), followed by Lun20 (SLUN_ RS38505, elongation module 3), and finally Lun22 (SLUN_RS38515, elongation and termination modules 4, 5, 6, and 7) (**Figure 1**). All together these NRPSs form a megasynthetase organized into seven modules, for which the detailed analysis of the specificity-conferring sequence of each adenylation (A) domains with SANDPUMA [39] allowed the prediction of the building blocks recruited by the NRPS machinery (**Figure 1**).

Module 1 in Lun23 contains three domains (A-T-E) and is predicted to incorporate L-Phe (DAWTVAAVCK as 10-aa code), which would be converted to D-Phe by the epimerization (E) domain. Module 2 in Lun23 also contains three domains (C-A-T) but prediction software tools did not propose specificity to any amino acid and suggest that this module would be inactive (DVASLAAYAK as 10-aa code). Module 3 in Lun20 contains three domains (C-A-T) and is predicted to recruit L-Ile (DAFFLGVTYK as 10-aa code). The four following modules (4 to 7), all included in Lun22, are predicted to incorporate four Piz residues, two D-Piz via modules 4 (C-A-T-E) and 6 (C-A-T-E) that both contain an epimerization domain, and two L-Piz via module 5 (C-A-T) and module 7 (C-A-T-TE). Indeed, the 10 aa code sequence conferring amino acid substrate specificity for Lun22 (DVFSVAAYAK as 10-aa code for modules 4, 5, 6, and 7) is identical to the one of HmtL-A1 (DVFSVAAYAK), and highly similar to those of KtzH-A1 (DVFSVGPYAK as 10-aa code, 8/10), ArtF-A1 (DVFSVASYAK as 10-aa code, 9/10), and ArtG-A2 (DVFTVAAYAK as 10-aa code, 9/10) which were reported to recognize and activate Piz in previous studies [25,28,30,37,38]. Finally, Module 7 would cyclize and release the final hexapeptide via its thioesterase (TE) domain as shown for the biosynthesis of himastatins and kutznerides where the last TE domain of HmtL and KtzH, the homologues of Lun22, perform the macrocyclization of their respective final peptide [25,28].

In conclusion, the NRPS machinery of BGC 28a is predicted to generate an hexapeptide with the sequence D-Phe, L-Ile, D-Piz, L-Piz, D-Piz, L-Piz as backbone of its main product that, with or without cyclization, would have monoisotopic masses of 708.4071 Da (C_35_H_52_N_10_O_6_) and 726.4177 Da (C_35_H_54_N_10_O_7_), respectively.

### 2.2. Structure elucidation of lunaemycins

#### 2.2.1 MS/MS-based networking and peptidogenomics guided genome mining

A peptidogenomic approach where genome mining guides the tandem mass spectrometry (MS/MS)-based molecular identification of high-resolution mass spectrometry (HRMS) data was used to identify compounds produced by BGC 28a. The predicted building-blocks can be used to screen the MS/MS fragments in order to find neutral or charged losses of amino-acids. Although BGC 28a is predicted to produce a cyclized hexapeptide of a monoisotopic mass of 708.4071 Da, we cannot exclude at this stage either modifications of one or more building blocks before their utilization by the NRPS machinery and/or post modifications of the generated hexapeptide. Therefore, the MS/MS spectra of molecular ions ranging from m/z 700 to 800 Da have been manually analyzed to identify tag fragments corresponding to the three different amino acids predicted to be incorporated by the six NRPS modules. Finally, prior to performing the comparative metabolomic analysis, genome mining of the eighteen isolated *S. lunaelactis* strains revealed a genetic characteristic that is crucial for the identification of the products of BGC 28a. Indeed, BGC 28a is present in only three *S. lunaelactis* strains i.e., those that possess pSLUN1a, whereas all the other strains with the linear plasmid pSLUN1b instead comprise BGC 28b [22] (**Figure 2a**). *S. lunaelactis* strains that possess pSLUN1b are therefore natural variants/mutants non-producing the searched compound(s) of BGC 28a and their culture extract can be considered as negative control.

**Figure 2.**
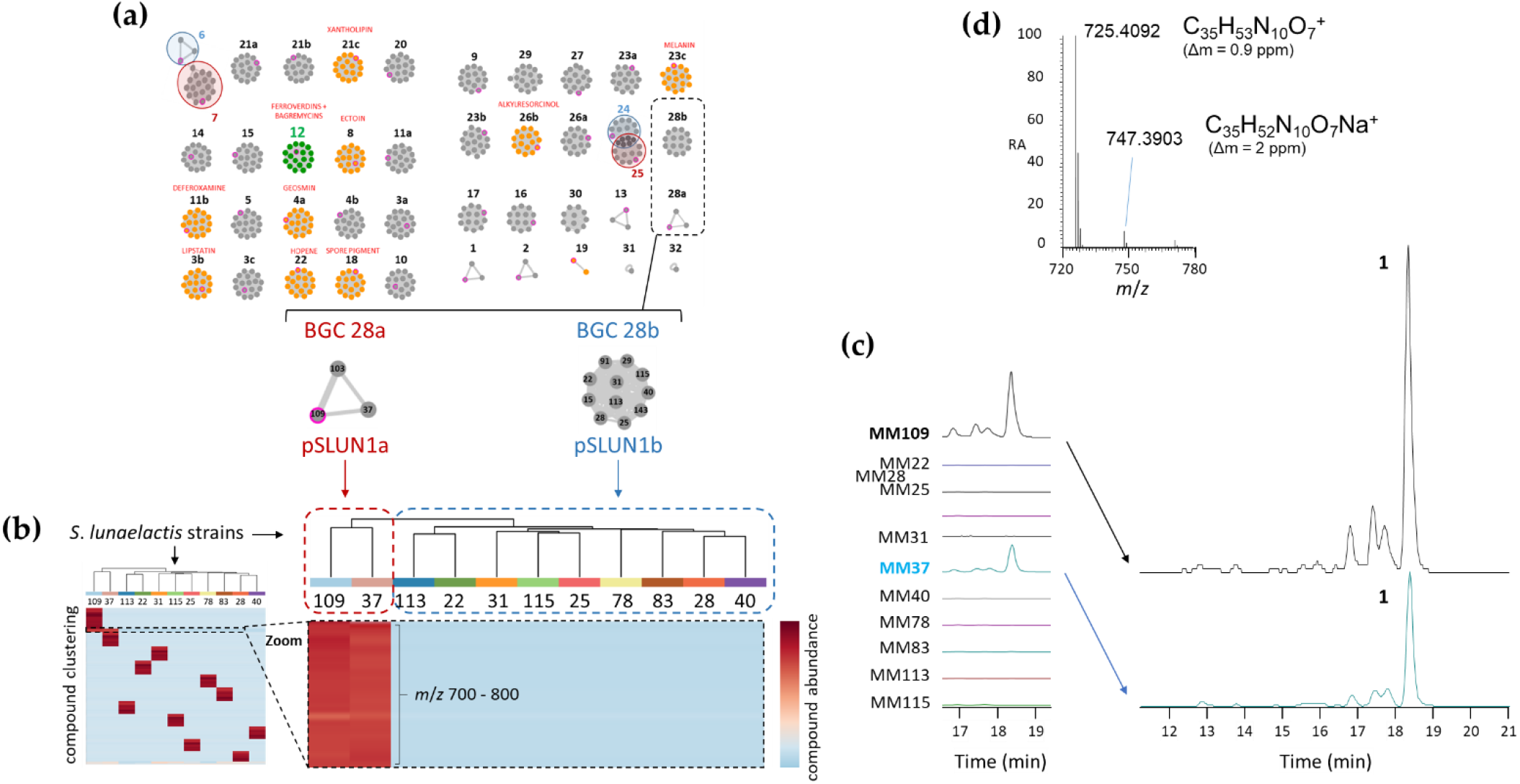
(**a**) Genome mining of *S. lunaelactis* strains where the 42 predicted BGCs are grouped in ‘grapes’ of nodes where each node represents a BGC in one strain (from [22]). Note the two grapes corresponding to BGC 28a and to BGC 28b found in strains either containing the linear plasmid pSLUN1a or pSLUN1b, respectively. (**b**) Heatmap of the metabolomic analysis of 11 *S. lunaelactis* strains grown on ISP7 media. The zoom on the heatmap highlights the differential production patterns for metabolites with *m/z* comprised between 700 and 800 Da and only present in the extract of *S. lunaelactis* strains that possess BGC 28a (MM109^T^ and MM37). (**c**) Extracted ion chromatograms (EIC, from *m/z* 700 to 800) of the full extract of the 11 strains with a focus on the retention times of signals associated with compounds only detected in *S. lunaelactis* strains with BGC 28a (MM109^T^ and MM37). (**d**) HRMS-predicted molecular formula of compound **1** (for both the proton and the sodium adducts) associated with the main signal detected in EICs of *S. lunaelactis* strains MM37 and MM109^T^.

The crude extracts from eleven *S. lunaelactis* strains grown on the ISP7 solid medium were collected and subjected to UPLC-HRMS/MS analysis. As explained above, metabolite profiling was performed to search for molecules with *m/z* comprised between 700 and 800 Da that were only produced by the two selected *S. lunaelactis* strains that possess pSLUN1a (MM109^T^ and MM37), and not by the nine other selected strains that instead have pSLUN1b (MM22, MM25, MM28, MM31, MM40, MM78, MM83, MM113 and MM115) (**Figure 2b**). As shown in the heatmap of **Figure 2b**, a series of *m/z* signals displayed contrasting production patterns between the two groups of *S. lunaelactis* strains. Analysis of extracted ion chromatograms (EIC) of the full extract of the 11 *S. lunaelactis* strains revealed a series of peaks only present in strains harboring pSLUN1a (**Figure 2c**). The most intense signal has an *m/z* of 725.4092 appropriate for the molecular formula of C_35_H_53_N_10_O_7_^+^ (0.17 ppm error) (**Figure 2d**). In the same HRMS spectrum a second ion with lower signal intensity corresponds to the [M + Na]^+^ ion at *m/z* 747.3903 which corresponds to the molecular formula C_35_H_52_N_10_O_7_Na^+^. The masses and formula of both proton and sodium adducts reveal that the main compound of BGC 28a has a molecular formula of C_35_H_52_N_10_O_7_ (monoisotopic mass of 724.4020 Da) from now on referred as compound **1** (hereafter named lunaemycin A).

Interestingly, the experimentally calculated mass of compound **1** only differs by 16 Da compared to the mass of the *in silico* predicted cyclized structure of the main product of BGC 28a (**Figure 1**). From the molecular formula C_35_H_53_N_10_O_7_ inferred from *m/z* 725.41, and according to the amino acid building blocks (D-*allo*-Ile, L-Phe, D-Piz, L-Piz, D-Piz, L-Piz) predicted to be loaded by adenylation domains, compound **1** could correspond to either i) an open peptide with an extra unsaturation from a dehydrogenation step, or ii) a cyclic peptide carrying a hydroxyl group added in an oxidation step.

Analysis of the HRMS/MS spectrum of the *m/z* 725.41 molecular ion revealed sequence tags that allowed us to propose that compound **1** is the cyclized sequence of the hexapeptide Phe-OH-Ile-Piz-Piz-Piz-Piz (**Figure 3**). Indeed, the daughter ion fragments of *m/z* 596.33, 449.26, 337.2, 225.13, and 113.07 correspond to consecutive losses of i) an hydroxy-leucine (H_2_O neutral loss-derived tag fragment of *m/z* 129.08), ii) a phenylalanine (H_2_O neutral loss-derived tag fragment of *m/z* 147.07), followed by three consecutive losses of Piz residues (three H_2_O neutral loss-derived tag fragments of *m/z* 112.06), and finally the ion fragment tag of *m/z* 113.07 corresponding to Piz+H^+^ (**Figure 3b**). The fragmentation spectrum thus confirmed the presence of tags associated with all six amino acids predicted to be incorporated by the A domains of the active modules of BGC 28a. The only difference with the initial prediction is the hydroxylation of the Ile residue which could be attributed to the product of *lun21* (SLUN_RS38510, **Table 1**) encoding a FAD-dependent oxidoreductase, homologous to the oxidoreductase PlyE of *Streptomyces* sp. MK498-98 F14 involved in the production of polyoxypeptins [40]. Indeed, PlyE is predicted to be responsible for the N-hydroxylation of valine and alanine building blocks on the nitrogen atom between Piz and Val or Ala residues of polyoxypeptin A. A similar role is also proposed for the FAD-dependent monooxygenase CchB that catalyzes the N-hydroxylation of the δ-amino group of ornithine in coelichelin biosynthesis before its recruitment by the NRPS machinery [41]. Similarly, Lun21 would perform the N-hydroxylation of the Ile residue, which would lead to the structure proposed for compound **1** (**Figure 3b**). Whether this N-hydroxylation of the Ile building block is performed i) before its recruitment by the A domain of Lun20 (as suggested for coelichelin biosynthesis), ii) once it is activated and tethered on thiolation (T) domain of Lun20 (as proposed for polyoxypeptin synthesis), or after formation and cyclization of the hexapeptide, remains speculative at this stage.

**Figure 3.**
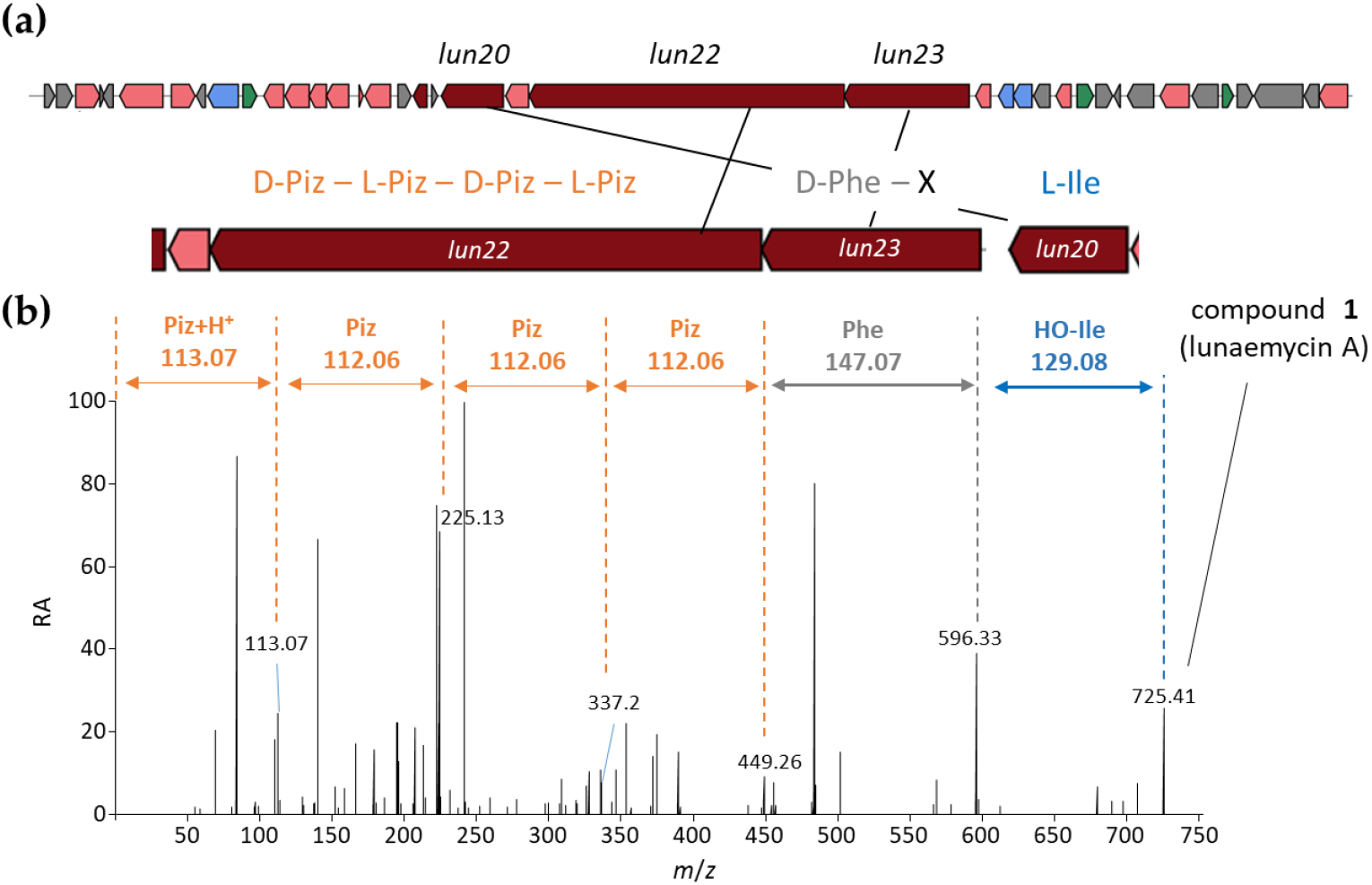
HRMS/MS-based identification of the amino acid building-blocks of compound **1**. (**a**) NRPS genes of BGC 28a (*lun* cluster) and the predicted amino acid building-block sequence. (**b**) MS fragmentation spectrum and proposed structure of compound **1** (lunaemycin A). Only fragments corresponding to amino-acid or peptide *m/z* of the predicted peptide sequence are highlighted. Color code: orange, piperazic acid (Piz); grey, phenylalanine (Phe); blue, hydroxylated isoleucine (HO-Ile).

To strengthen the reliability of the proposed structure of compound **1**, the *m/z* of the MS/MS data were correlated to the structure of ions produced during the fragmentation mechanism (**Figure 4a**). The *m/z* of nine fragment ions can only correlate with the structure of ions containing one to four Piz residues (fragments 1-9 in blue in **Figure 4b**). Four of these fragment ions – *m/z* 113.0711, 225.1347, 337.1984, and 449.2608 – can be linked to structures containing respectively 1, 2, 3, and 4 Piz residues, expected to result from sequential *b_x_*–*y_z_* fragmentation steps. Additionally, *m/z* 309.2034 and 197.1394 result from the neutral loss of CO of the aforementioned ions *m/z* 337.1984 and 225.1347, respectively. At least seven fragment ions contain phenylalanine’s aromatic side chain thereby confirming the presence of this amino acid in the structure of compound **1** (fragments 10-16 in green in **Figure 4c**). Fragment ions with *m/z* 260.1396, 372.2032, 484.2668, and 596.3304, arise from sequential *b_x_*–*y_z_* fragmentation steps, and represent phenylalanine connected to increasing numbers of piperazate residues, whereas the remaining ions (*m/z* 232.1445, 456.2719, and 568.3356) are associated to a subsequent CO loss step.

**Figure 4.**
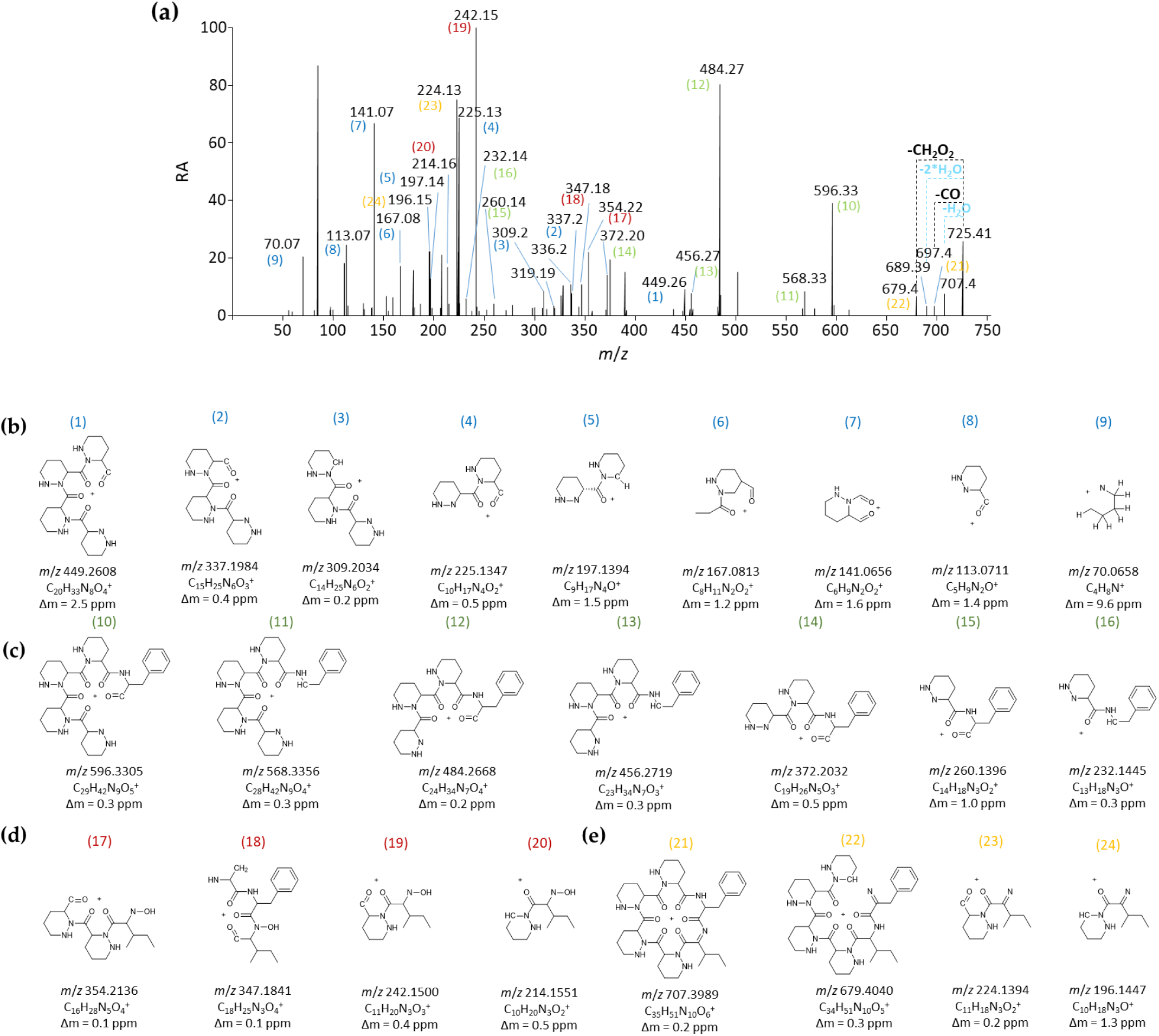
HRMS/MS-based identification of fragmentation patterns of compound **1**. (**a**) MS fragmentation spectrum. (**b**) Structure of ions containing one to four Piz residues (these ions are highlighted by numbers in blue in the MS fragmentation spectrum). (**c**) Structure of fragment ions containing the phenylalanine building block (these ions are highlighted by numbers in green in the MS fragmentation spectrum. (**d**) Structure of fragment ions that contain a hydroxyl-isoleucine residue (these ions are highlighted by numbers in red in the MS fragmentation spectrum). (**e**) Structure of fragment ions seemingly resulting from the neutral loss of H_2_O further supporting the presence of hydroxylated isoleucine (these ions are highlighted by numbers in orange in the MS fragmentation spectrum).

It is interesting to note that no hydroxylation can be inferred from the seventeen fragments discussed above, confirming that the hydroxylation in compound **1** must reside in the remaining amino acid, i.e., isoleucine. This is confirmed by the presence of four fragments that contain a hydroxyl-isoleucine residue (*m/z* 214.1551, 242.1500, 347.1841 and 354.21366, fragments 17-20 in red in **Figure 4d**), all of which resulting from similar fragmentation steps as discussed for the other ions (sequential *b_x_*–*y_z_* fragmentation and CO loss). Finally, the hydroxylation of compound **1** and the hypothesis of a hydroxylated isoleucine in its structure are further confirmed by the presence of four fragment ions seemingly resulting from the neutral loss of H_2_O (fragments 21-24 in orange in **Figure 4e**). The fragment ion with *m/z* 707.3989 is the result of a water loss from the [M+H]^+^ adduct and can itself generate the ion with *m/z* 679.4040 from a CO loss. The two other fragment ions (*m/z* 196.1447 and 224.1394) would come from the loss of water from previously mentioned isoleucine-bearing fragments *m/z* 214.1551 and 242.1500, respectively.

#### 2.2.2 Structure elucidation by NMR spectroscopy

Finally, nuclear magnetic resonance (NMR) was performed in order to confirm the predicted and MS/MS-deduced structure of compound **1**. Indeed, even though we can confidently propose the presence of a hydroxy-leucine residue in the structure of compound **1**, the MS/MS information alone does not allow to specify the position of the hydroxyl group in this residue. The fragment structures proposed in **Figure 4d** assume a hydroxylation on the amide nitrogen of isoleucine, which can only be confirmed by NMR. Multiple media were tested (OSMAC approach [42] and known elicitors of secondary metabolite production in *Streptomyces* spp. [43–45]) to assess the optimal culture conditions for compound **1** production in order to obtain sufficient material for NMR analysis. Analysis of the metabolite profiles showed that the solid ISP1 medium supplemented with N-acetyl-D-glucosamine 25 mM led to the best production yield of compound **1** (not shown). The dried ethyl-acetate full extract of *S. lunaelactis* MM109^T^ was processed using the ÄKTA liquid chromatography purification system, followed by semi-preparative HPLC (SP-HPLC) purification, which led to 2.15 mg of a pale-yellow-white powder from ~0.5 liters of solid culture. UPLC-HRMS/MS revealed that this fraction contained two compounds with similar molecular weights, the most intense molecular ion peak being compound **1** (*m/z* of 725.4092), and a second molecular ion peak with an *m/z* of 723.3935 (compound **2**). This *m/z* value suggests that compound **2** is a dehydrogenated derivative of compound **1**, possibly containing a dehy-dro-Piz moiety as described previously for antrimycins [46]. This hypothesis was confirmed by ^1^H NMR results (see below), and by HRMS/MS analysis (see the following section on the structural diversity of lunaemycins).

The semi-purified powder containing compound **1** and minor amount of compound **2** was submitted to ^1^H NMR (700MHz), ^13^C NMR (176 MHz) as well as 2D NMR (COSY, HMBC, and HSQC). 1D- and 2D NMR data were analyzed with MestReNova V.14 and allowed to confirm the chemical structure predicted by the *in silico* analysis of BGC 28a and deduced from the MS/MS fragmentation pattern. **Table 2** shows the structure and NMR assignment proposed for compounds **1** and **2** (**Figure 5a** and **5b** and **supplementary Figures S2-S6**), however, the overlapping signals did not allow a complete structural assignment, as multiplicities and integrations were not unambiguously characterized.

**Figure 5.**
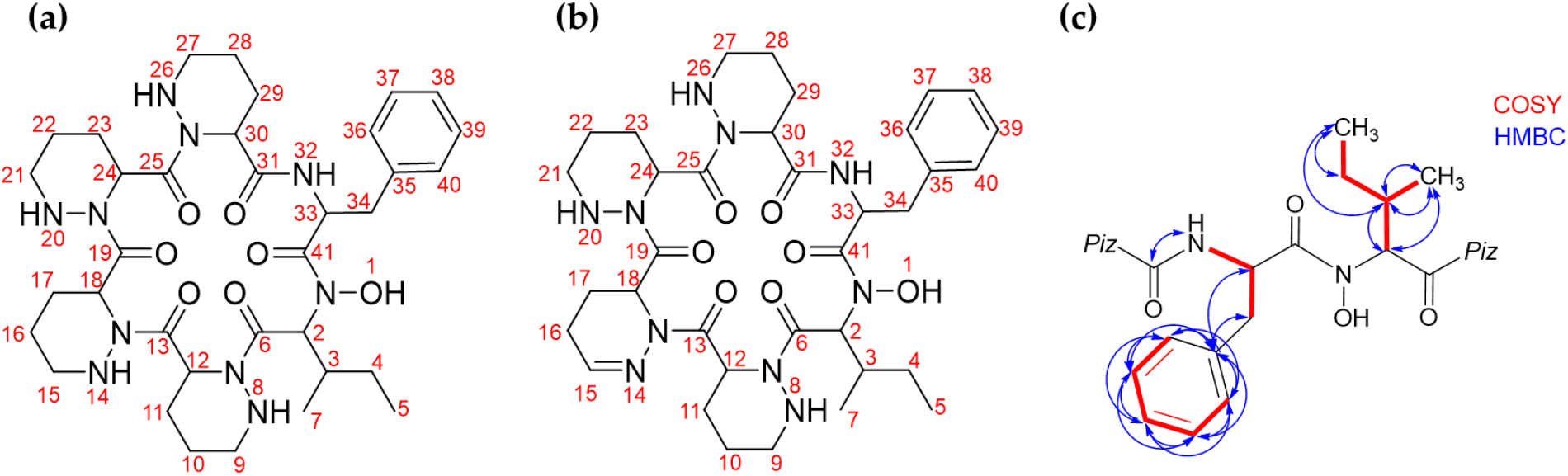
Structure elucidation of compounds 1 and 2 by NMR. (**a**) Compound **1**. (**b**) Compound **2**. (**c**) COSY and HMBC correlations that allowed to unambiguously confirm the location of the hydroxylation on compound 1. See supplementary **Figures S2-S6**.

**Table 2.** Assignment of NMR signals of the major (1) and minor (2) components of the semi purified sample.

| atom # | Major compound (1) |  | Minor compound (2) |  |
| --- | --- | --- | --- | --- |
|  | <sup>1</sup>H NMR<sup>a</sup> | <sup>13</sup>C NMR<sup>b</sup> | <sup>1</sup>H NMR<sup>a</sup> | <sup>13</sup>C NMR<sup>b</sup> |
| 1 | 9.27 (b, 1H) |  | 9.64 (b, 1H) |  |
| 2 | 5.82-5.76 | 56.4 | 5.96 (d, $J = 11.0$ Hz, 1H) | 55.6 |
| 3 | 2.07 _ 2.03 | 32.8 | 2.12 _ 2.07 | 32.3 |
| 4 | 1.02-0.94 | 25.2 | 1.02-0.94 | 25.2 |
| 5 | 0.82-0.76 | 11.1 | 0.82-0.76 | 10.9 |
| 6 |  | 172.7 |  | 172.2 |
| 7 | 0.88-0.84 | 15.5 | 0.88-0.84 | 15.4 |
| 8 | 4.92-4.84<sup>c</sup> |  | 4.92-4.84<sup>c</sup> |  |
| 9 | 2.9-3.1<sup>d</sup> | 46.9<sup>d</sup> | <sup>e</sup> | <sup>e</sup> |
| 10 | 1.2-2.3<sup>f</sup> | 18.4-26.2<sup>f</sup> | <sup>e</sup> | <sup>e</sup> |
| 11 | 1.2-2.3<sup>f</sup> | 18.4-26.2<sup>f</sup> | <sup>e</sup> | <sup>e</sup> |
| 12 | 5.75 (d, $J = 5.6$ Hz, 1H)<sup>g</sup> | 48.9<sup>g</sup> | 5.82- 5.76<sup>g</sup> | 47.6<sup>g</sup> |
| 13 |  | 173.5<sup>h</sup> |  | 171.3<sup>h</sup> |
| 14 | 5.05 (d, $J = 13.8$ Hz, 1H)<sup>c</sup> | | | |
| 15 | 2.9-3.1<sup>d</sup> | 47.1<sup>d</sup> | 6.91 (d, J = 3.7 Hz, 1H)<sup>i</sup> | 142.9<sup>i</sup> |
| 16 | 1.2-2.3<sup>f</sup> | 18.4-26.2<sup>f</sup> | <sup>e</sup> | <sup>e</sup> |
| 17 | 1.2-2.3<sup>f</sup> | 18.4-26.2<sup>f</sup> | <sup>e</sup> | <sup>e</sup> |
| 18 | 5.39<sup>g</sup> | 48.1<sup>g</sup> | 5.36-5.28<sup>g</sup> | 47.5<sup>g</sup> |
| 19 |  | 174.1<sup>h</sup> |  | 172.9<sup>h</sup> |
| 20 | 5.18 (d, J = 13.0 Hz, 1H)<sup>c</sup> |  | 5.01 (d, J = 14.4 Hz, 1H)<sup>c</sup> |  |
| 21 | 2.9-3.1<sup>d</sup> | 47.2<sup>d</sup> | <sup>e</sup> | <sup>e</sup> |
| 22 | 1.2-2.3<sup>f</sup> | 18.4-26.2<sup>f</sup> | <sup>e</sup> | <sup>e</sup> |
| 23 | 1.2-2.3<sup>f</sup> | 18.4-26.2<sup>f</sup> | <sup>e</sup> | <sup>e</sup> |
| 24 | 5.45<sup>g</sup> | 47.6<sup>g</sup> | 5.67<sup>g</sup> | 46.7<sup>g</sup> |
| 25 |  | 173.2<sup>h</sup> |  | 171.6<sup>h</sup> |
| 26 | 5.23 (dd, J = 12.8, 3.4 Hz, 1H)<sup>c</sup> |  | 5.36-5.28<sup>c</sup> |  |
| 27 | 2.9-3.1<sup>d</sup> | 47.3<sup>d</sup> | <sup>e</sup> | <sup>e</sup> |
| 28 | 1.2-2.3<sup>f</sup> | 18.4-26.2<sup>f</sup> | <sup>e</sup> | <sup>e</sup> |
| 29 | 1.2-2.3<sup>f</sup> | 18.4-26.2<sup>f</sup> | <sup>e</sup> | <sup>e</sup> |
| 30 | 4.92-4.84<sup>g</sup> | 49.6<sup>g</sup> | 5.37<sup>g</sup> | 47.6<sup>g</sup> |
| 31 |  | 170.9(8)<sup>h</sup> |  | 171.0(2)<sup>h</sup> |
| 32 | 8.27 (d, J = 8.6 Hz, 1H) |  | 8.22 (d, J = 9.0 Hz, 1H) |  |
| 33 | 5.36-5.28 | 49.0 | 5.68 (d, J = 5.4 Hz, 1H) | 49.1 |
| 34 | 2.93-2.86 and 2.77-2.67 | 37.7 | 2.93-2.86 and 2.77-2.67 | 37.7 |
| 35 |  | 137.7(1) |  | 137.7(4) |
| 36 | 7.20-7.14<sup>j</sup> | 129.7(0) | 7.20-7.14<sup>j</sup> | 129.7(3) |
| 37 | 7.24<sup>j</sup> | 128.4 | 7.24<sup>j</sup> | 128.3 |
| 38 | 7.20-7.14<sup>j</sup> | 126.6(7) | 7.20-7.14<sup>j</sup> | 126.6(5) |
| 39 | 7.24<sup>j</sup> | 128.4 | 7.24<sup>j</sup> | 128.3 |
| 40 | 7.20-7.14<sup>j</sup> | 129.7(0) | 7.20-7.14<sup>j</sup> | 129.7(3) |
| 41 |  | 170.3 |  | 170.4 |

Considering the chemical shifts observed in the ^1^H and ^13^C NMR spectra, signature signals typical of cyclopeptides were observed revealing peptide bonds between amino acid residues with no amino or carboxy interruption of the amino acid sequence. ^1^H NMR data revealed the presence of the six α hydrogens of the hexapeptide core between δ 4.8 and δ 5.8 that can be attributed to compound **1** (δ 5.82-5.76 for the Ile; δ 5.36-5.28 for the Phe; δ 4.92-4.84, 5.39, 5.45, and 5.75 for the Piz residues), and compound **2** (δ 5.96 for the Ile; δ 5.68 for the Phe; δ 4.92-4.84, 5.01, 5.36-5.28, and 5.67 for the Piz residues). Further confirmation comes from the ^13^C-NMR signals between δ 170.3 and δ 174.1, attributed to the peptide carbonyl groups (C-6, C-13, C-19, C-25, C-31, C-41), and between δ 46.7 and δ 56.4, attributed to the α carbons (C-2, C-12, C-18, C-24, C-30, C-33).

Based on previously published data [47], a series of multiplets between δ 1.2 and δ 2.3 in the ^1^H NMR spectra can be attributed the eight β- and γ-methylene groups from the piperazate residues present in the structure of compound **1** (C-10, C-11, C-16, C-17, C-22, C-23, C-28, and C-29). Additionally, a series of ^13^C NMR signals between δ 18.4 and δ 26.2 can be attributed to those same methylene groups. Similarly, the four δ-methylene groups (C-9, C-15, C-21, and C-27) can be correlated to the ^1^H NMR signals between δ 2.9 and δ 3.1 and to the ^13^C NMR signals between δ 46.9 and δ 47.3. Interestingly, a signal at δ 6.91 (d, J = 3.7 Hz, ^1^H for compound **2**) in the ^1^H NMR spectra, which is correlated on the HSQC to the ^13^C NMR signal at δ 142.9, can be attributed to a methylene hydrogen of an imine group. This observation supports the hypothesis of a replacement of a piperazate residue by a dehydro-piperazate in compound **2**.

The presence of phenylalanine and isoleucine residues in the structure of compounds **1** and **2** is clearly detectable from the results of NMR experiments (**Table 2**). The six aromatic carbons (C-35 to C-40) of the phenylalanine’s benzene ring were observed at δ 137.7 (quaternary carbon, C-35); δ 128.4 (two meta carbons, C-37 and C-39); 129.7, (two ortho carbons, C-36 and C-40); and δ 126.6 (the para carbon, C-38). Also, the five aromatic hydrogens corresponding to this benzene ring were observed in ^1^H NMR spectra between δ 7.1 and δ 7.2. The two hydrogens at meta position (linked to C-37 and C-39) were attributed to the signal at δ 7.24, and could be discerned from the three hydrogens on ortho (linked to C-36 and C-40) and para (linked to C-38), seen at δ 7.2-7.14. As for the non-aromatic carbons in those residues, we were able to attribute ^13^C signals to the two methylenes (δ 25.2 for C-4 from the Ile and δ 37.7 for C-34 from the Phe), to the tertiary carbon C-3 from the Ile (δ 32.8), and to the two methyls, C-5 and C-7, from the Ile (δ 11.1 and 15.5, respectively). The upfield region of the ^1^H NMR spectrum shows signals corresponding to the non-aromatic hydrogens of the minor (**2**) and major (**1**) compounds of the sample analyzed by NMR. The methylene hydrogen of isoleucine, linked to C-4, can be attribute to δ 1.02-0.94, while the methylene hydrogens of phenylalanine, linked to C-34, were attributed to δ 2.93-2.86 and 2.77-2.67. For the isoleucine, the hydrogen on C-3 was attributed to peaks between δ 2.12-2.03, and the methyl hydrogens linked to C-5 and C-7 were attribute to δ 0.82-0.76 and δ 0.84, respectively.

The observation of the non-substituted s-butyl group of isoleucine indicates that hydroxylation of **1** is not localized at its side chain, suggesting that the hydroxylation should take place at the amide nitrogen. Additionally, 2D NMR COSY and ^1^H-^13^C HMBC correlations, shown on **Figure 5c**, further supported this conclusion. Another key observation is the presence of a single HMBC correlation between an alpha NH ^1^H NMR signal at δ 8.27 (attributed to phenylalanine’s NH) and an alpha carbonyl ^13^C NMR signal at δ 172.7 (attributed to the neighboring piperazate carbonyl). Those observations allowed us to further confirm the N-hydroxylated structure previously proposed for Lunaemycin A.

In conclusion, combined *in silico*, HRMS/MS, and NMR analyses demonstrated that the main product of BGC 28a, compound **1**, is a cyclic hexapeptide of a mass of 724.4019 Da and the molecular formula C_35_H_52_N_10_O_7_, corresponding to the cyclized amino acid sequence D-Phe-OH-L-Ile-D-Piz-L-Piz-D-Piz-L-Piz. We have interrogated StreptomeDB 3.0 [48] and The Natural Products Atlas 2.0 [49] databases and found no natural compounds with neither identical molecular formula, monoisotopic mass, nor molecular weight. Interrogation of databases with broader chemical compounds (NIST, Pubchem, and Chemspider) identified ‘SCHEMBL12568990’ (Substance SID: 237623651; Compound CID: 53234730 in the Pubchem database) retrieved from patent US-2011136752-A1 [50], as the closest match of compound **1**, sharing the same mass, same molecular formula, and same proposed structure. We decided to call compound **1** lunaemycin A, the Latin prefix lunae-(L. gen. n. lunae, of the moon) referring to the moonmilk speleothems where *S. lunaelactis* MM109^T^ was originally isolated, and the suffix-mycin which refers to antibiotic compounds derived from bacteria with fungus-like structure such as *Streptomyces*. The antibiotic activity of lunaemycins A and lunaemycin-related compounds will be described in section 2.4.

### 2.3. Structural diversity of lunaemycins

The comparative metabolomic study of metabolites with masses ranging between 700 and 800 Da (**Figure 2b**) suggested a broad diversity of lunaemycin-derived molecules produced by *S. lunaelactis* strains containing pSLUN1a. Using the MS/MS data we explored this diversity via global natural products social molecular networking (GNPS) as performed previously [22]. One constellation comprises 42 nodes associated with compounds only produced by the two selected *S. lunaelactis* strains that possess BGC 28a (MM37 and MM109^T^) and includes the ion at *m/z* 725 corresponding lunaemycin A (**Figure 6**).

**Figure 6.**
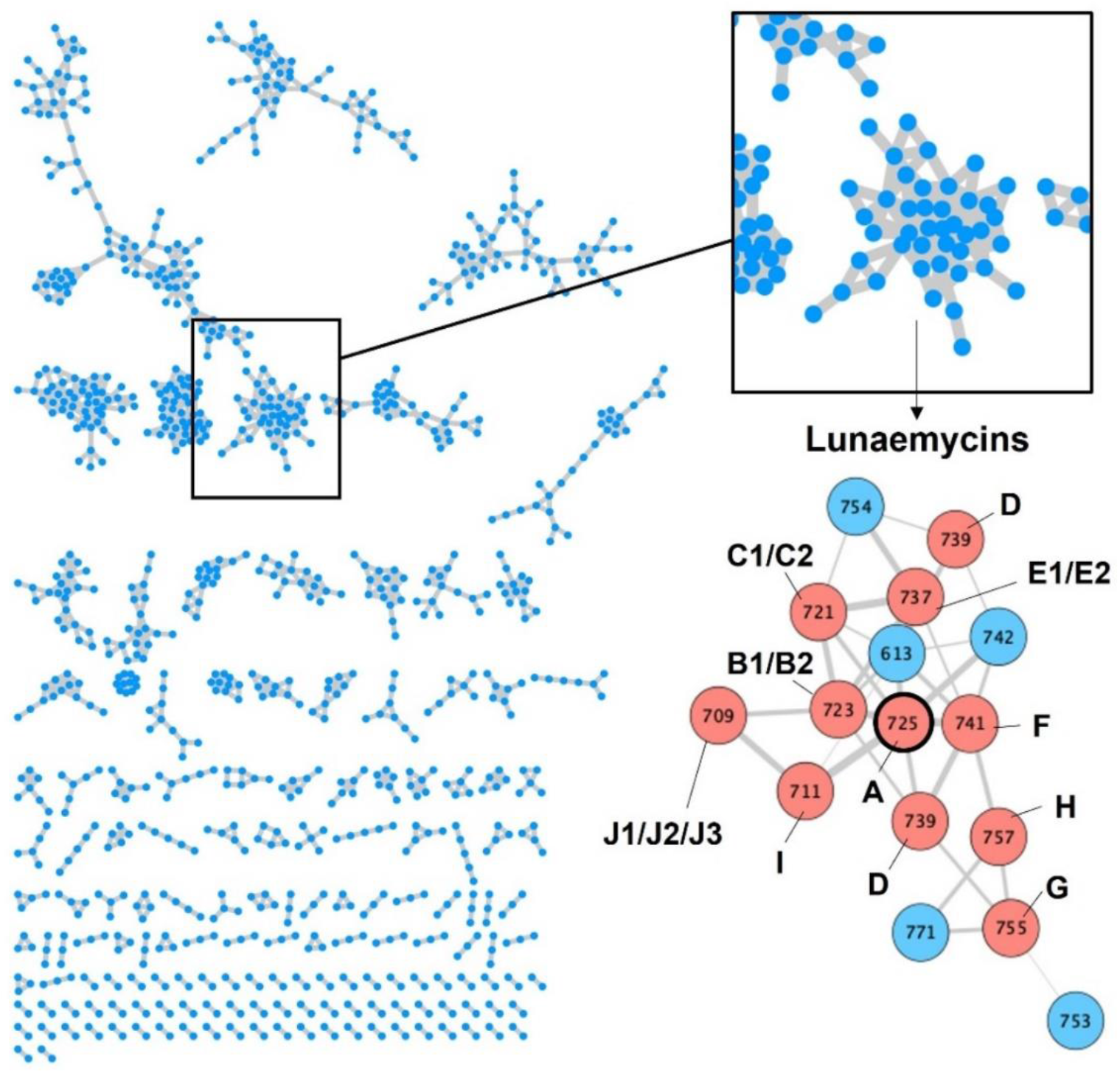
GNPS of metabolites extracted from *S. lunaelactis* strains. Left panel: Constellations of compounds constructed based on GNPS processing the UPLC-MS/MS data obtained from the full extract of the eleven *S. lunaelactis* strains grown on ISP7 medium. Each node in a constellation represents a parent mass (MS1) of a compound detected in one or more of the eleven full extracts analyzed. Right panel: Constellation of lunaemycins where nodes are linked by a straight line when they share similar fragmentation patterns. Pink and blue nodes: lunaemycins with solved and unsolved structure, respectively (see details in **Table 3, Figure 7** and supplementary Figure S7-S20)

**Figure 7** presents the detection of the most abundant compounds identified as lunaemycin derivatives produced by *S. lunaelactis* MM109^T^, as well as their HRMS spectra and their deduced molecular formula. Comparison of the MS/MS spectra allowed to identify the position of the chemical modification(s) for 14 of these molecules compared to lunaemycin A (Figure S7-S20, **Table 3**), whereas other five lunaemycin derivatives were produced in too weak amounts for proper analysis. **Table 3** lists all compounds clustering in the constellation of lunaemycins, their proposed name, retention time, *m/z* signals, and their molecular formula.

**Figure 7.**
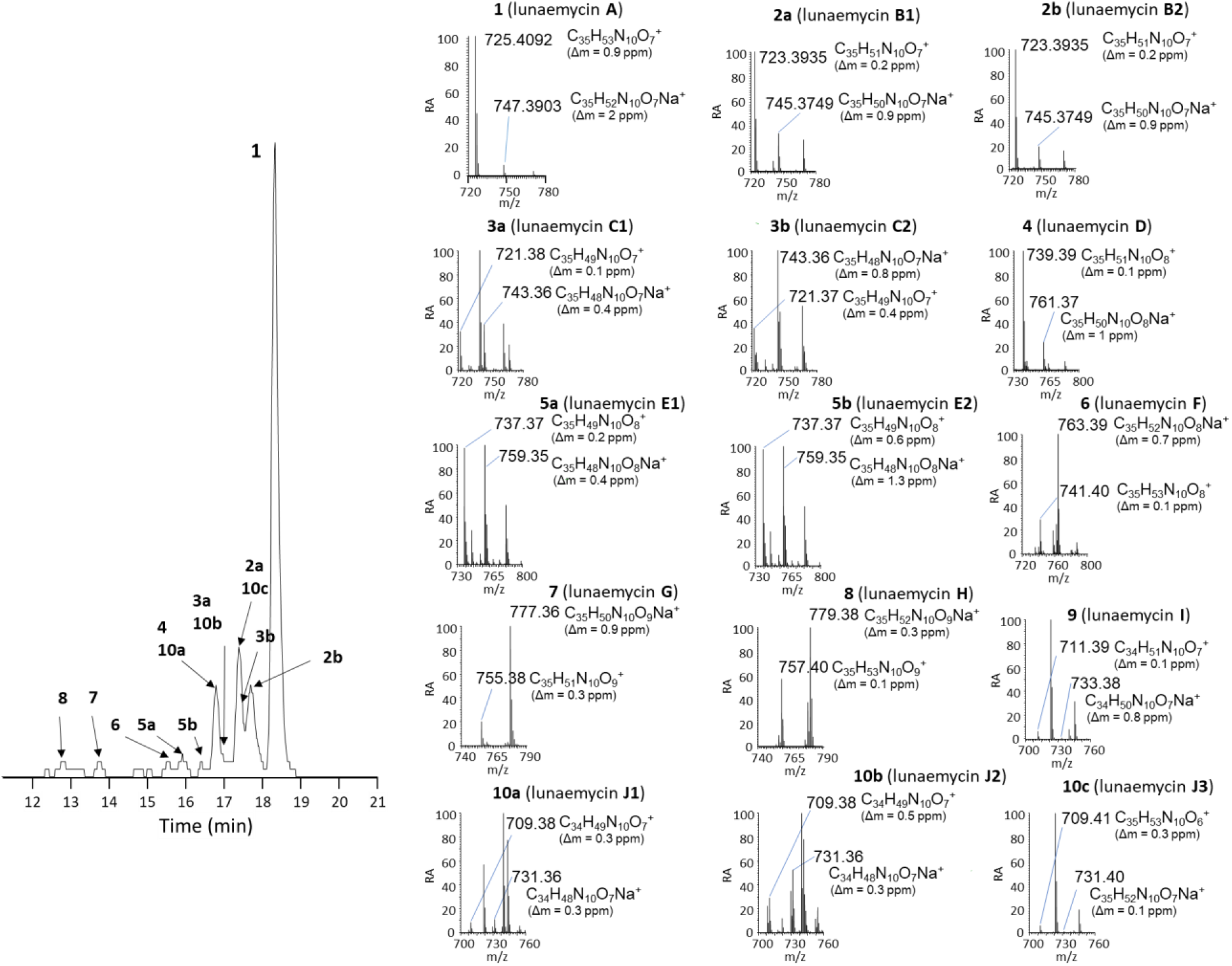
UPLC-HRMS chromatograms and mass spectra of lunaemycins. Left panel: Extracted ion chromatogram (EIC) with *m/z* signals ranging from 700 to 800 Da from the full extracts of *S. lunaelactis* MM109^T^ grown on ISP7 medium. Right panels: HRMS-predicted molecular formula of compound **1** to **10c** (for both the proton and the sodium adducts) associated with the main signal detected in the EIC displayed on the left panel.

**Table 3.**
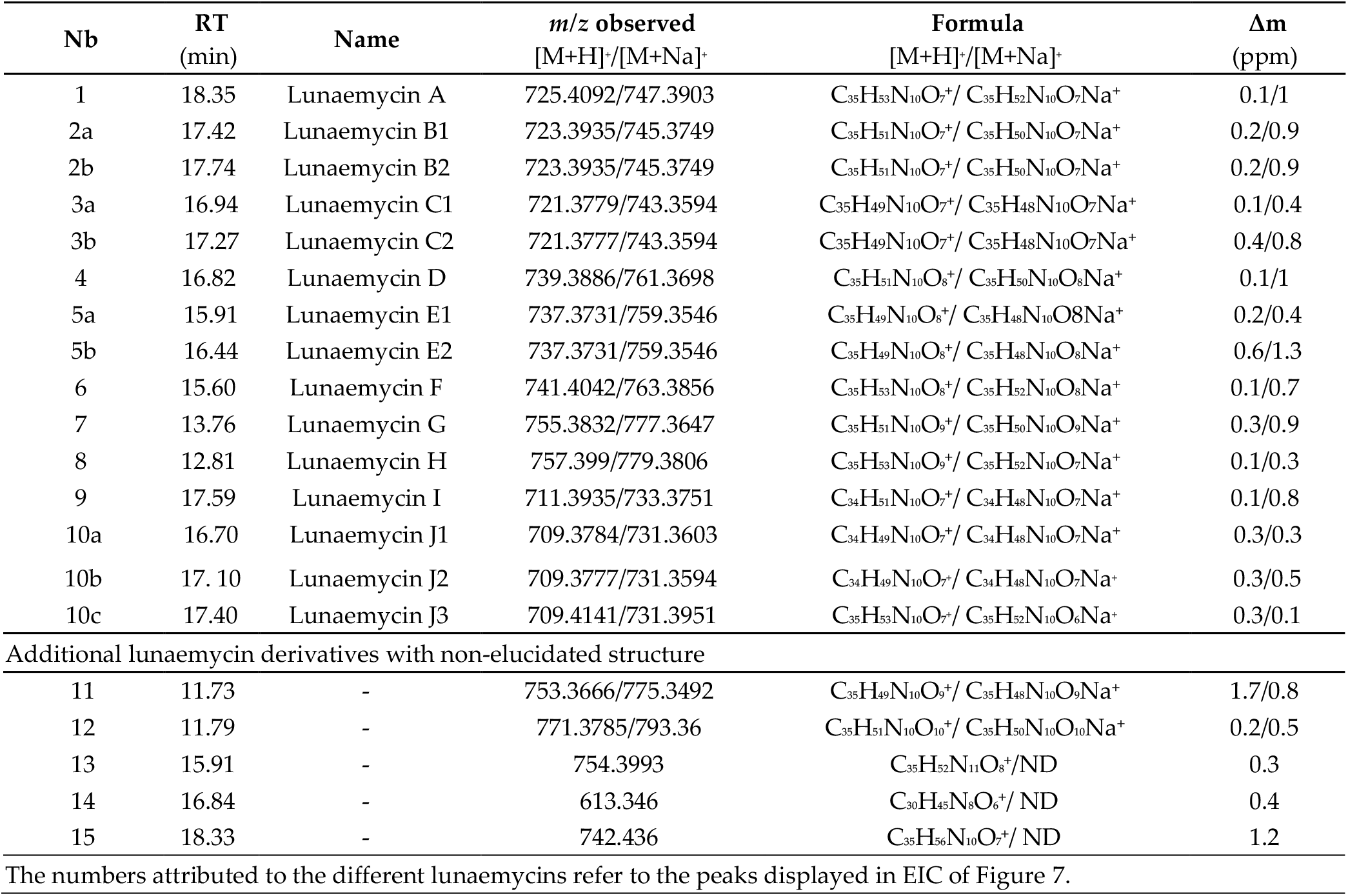
Lunaemycins produced by *S. lunaelactis* MM109^T^ and MM37.

The HRMS/MS-based identification of the amino acid building-blocks composing the 15 lunaemycins is detailed in supplementary **Figure S7-S20**. Eight lunaemycins possess at least one dehydrogenated-Piz (dh-Piz, corresponding to the neutral loss of 110.5) instead of a classical Piz. More precisely, if we consider the linear sequence of Lunaemycin A as Hydroxy-Ile-Piz1-Piz2-Piz3-Piz4-Phe, Piz1 is replaced by a dh-Piz in lunaemycin E**1** (**5a**); Piz2 is replaced by a dh-Piz in lunaemycins B**1** (**2a**), C**1** (**3a**), and J**1** (**10a**); Piz3 is replaced by a dh-Piz in lunaemycins B1(**2a**), C1 (**3a**), C2 (**3b**), E1 (**5a**), and J3 (**10c**); and finally, Piz4 is replaced by a dh-Piz in lunaemycin C1 (**3a**). Only lunaemycin F (**6**) presents the hydroxy-Piz residue (HO-Piz, corresponding to the neutral loss of 128.09) at the position of Piz3. Five lunaemycins present the hydroxydehydro-Piz residue (HO-dh-Piz, corresponding to the neutral loss of 126.05) at the position of Piz4 (lunaemycins D (**4**), E1 (**5a**), E2 (**5b**), G (**7**), and H (**8**). Finally, the HO-Ile is replaced by a dihydroxy-Ile (DIHO-Ile, corresponding to the neutral loss of 145.08) in lunaemycins G (**7**) and H (**8**), by an hydroxyl-Valine (HO-Val, corresponding to the neutral loss of 115.06) in lunaemycins I (**9**), J2 (**10b**), and J3 (**10c**), and by an Ile (corresponding to the neutral loss of 113.08) in lunaemycin J1 (**10a**).

### 2.4. Antibacterial activity of lunaemycins

Molecules produced by known BGCs closest to the *lun* cluster (himastatins, stenothricins, kutznerides, and aurantimycins) possess antimicrobial activities suggesting that lunaemycins could possibly display similar biological activities. Moreover, amongst the genes of the *lun* BGC, BLAST analyses also suggest that lunaemycins are likely to possess antibacterial and/or anti-proliferative activities. For instance, *lun9* and *lun10* encode for an efflux pump and its associated TetR-family transcriptional regulator, respectively, and whose homologues in other BGCs play a role in self-resistance of the producing organism. In addition, *lun25* and *lun26* and their closest homologues in the BGCs of aurantimycins (ArtJ and ArtK) and the polyoxipeptins (PlyJ and PlyK), encode for an ABC transporter system belonging to the DrrA-DrrB family also conferring self-resistance to the producers [51].

Antibacterial activity of lunaemycins A, B1, and D (0.1 mg/ml, ~13.5-14 mM) was first performed via agar-diffusion assays which showed significant direct-acting activity on all tested Gram-positive bacteria except on *Enterococcus hirae* (though a small zone of growth inhibition is observed at the periphery of the disc suggesting more susceptibility to lunaeemycins compared to gentamycin 0.1 mg/ml used as positive control) (**Figure 8**). In contrast, no growth inhibition was reported on Gram-negative bacteria (**Figure 8**). Overall, similar antibacterial activities could be observed for lunaemycins A and B1, whereas lunaemycin D showed weaker growth inhibiting activities.

**Figure 8.**
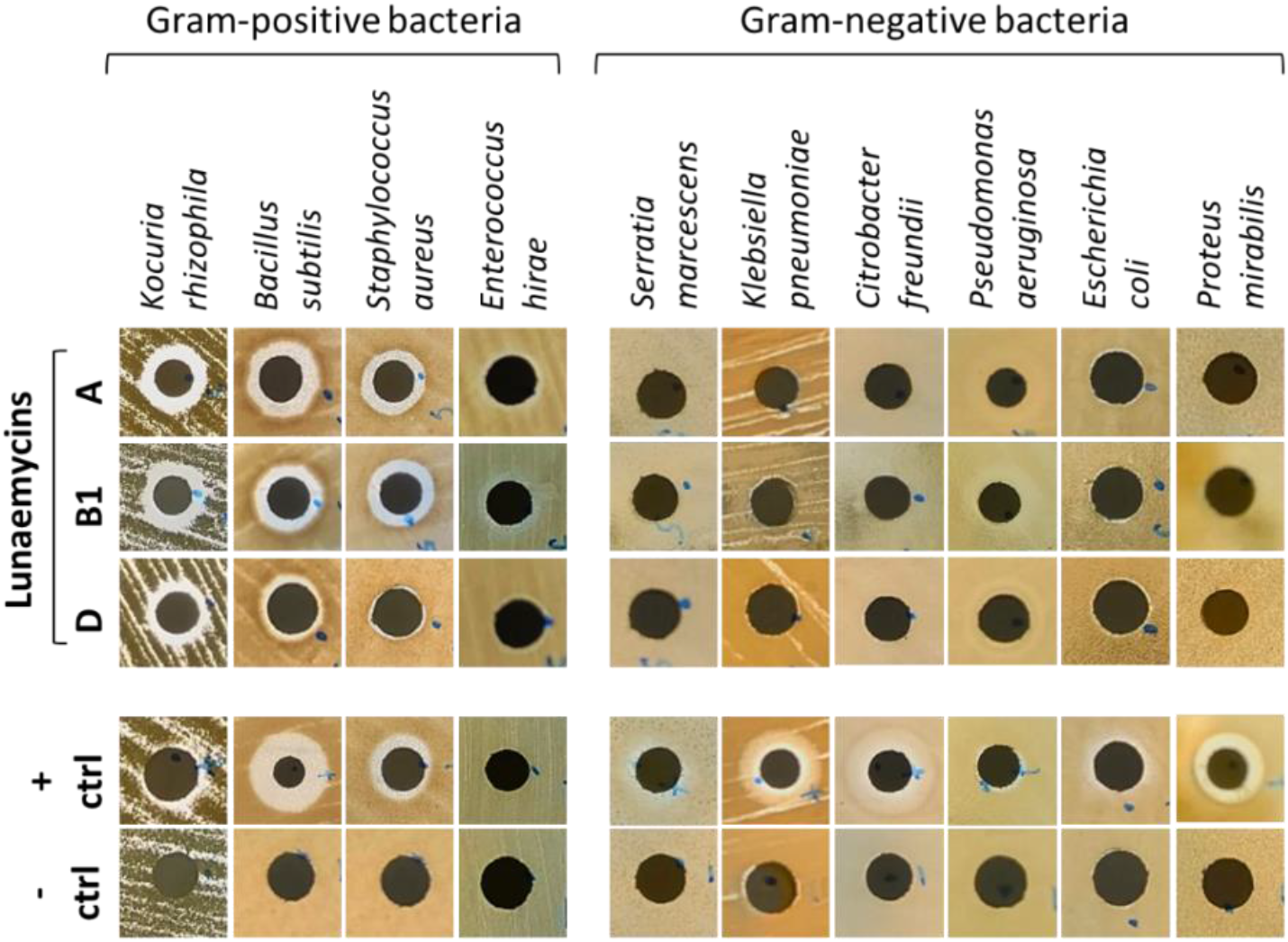
Agar-diffusion antibacterial activity assays of lunaemycins. 0.1 mg/ml of gentamycin was used as positive control, and a water/acetonitrile solution (1/9, v/v) was used as negative control.

Minimal inhibitory concentration (MIC) and minimal bactericidal concentration (MBC) values were determined with pure lunaemycin A. The potent direct-acting activity observed in agar diffusion assays was confirmed, with MIC values ranging 0.12-0.25 μg/ml on type strains and, more interestingly, also on clinical isolates showing various resistance phenotypes (**Table 4**). The MBC values were comprised between 0.25 and 8 μg/ml. Interestingly, the MBC values obtained with some antibiotic resistant clinical isolates, especially those showing resistance to methicillin, vancomycin or linezolid, were overall lower (from 0.25 to 2 μg/ml) than that of other strains, indicating a potent bactericidal activity of luneamycin A for these strains. Expectedly, no activity could be observed on both susceptible and resistant Gram-negative isolates as previously observed via the agar-diffusion assays.

**Table 4.** *In vitro* antibacterial activity (MIC and MBC values) of lunaemycin A on reference strains and clinical isolates with various phenotypes of resistance to antibiotics.

| Organism | Resistance | MIC (µg/ml) | MBC (µg/ml) |
| --- | --- | --- | --- |
| **Gram-positive bacteria** |  |  |  |
| *Bacillus subtilis* ATCC 6633 | - | 0.12 | 4 |
| *Enterococcus faecalis* ATCC 29212 | - | 0.12 | 8 |
| *Streptococcus pyogenes* ATCC 12344 | - | 0.12 | 8 |
| *Staphylococcus aureus* ATCC 25923 | - | 0.12 | 4 |
| *Enterococcus faecalis* SI-759 | vancomycin<sup>R</sup>, teicoplanin<sup>R</sup> | 0.25 | 8 |
| *Enterococcus faecium* SI-1831 | vancomycin<sup>R</sup> | 0.25 | 0.25 |
| *Staphylococcus aureus* ATCC 43300 | methicillin<sup>R</sup>, MRSA | 0.12 | 2 |
| *Staphylococcus epidermidis* SI-1266 | linezolid<sup>R</sup> | 0.12 | 0.25 |
| *Staphylococcus haemolyticus* SI-6/2011 | oxacillin<sup>R</sup> | 0.12 | 2 |
| *Staphylococcus warneri* SI-5/2011 | oxacillin<sup>R</sup> | 0.12 | 8 |
| **Gram-negative bacteria** |  |  |  |
| *Acinetobacter baumannii* ATCC 17978 | - | >64 | - |
| *Escherichia coli* CCUG<sup>T</sup> | - | >64 | - |
| *Klebsiella pneumoniae* ATCC 13833 | - | >64 | - |
| *Pseudomonas aeruginosa* ATCC 27853 | - | >64 | - |
| *Acinetobacter baumannii* N50 | pandrug-resistant | >64 | - |
| *Escherichia coli* SI-63 | MDR, colistin<sup>R</sup>, mcr<sup>+</sup> | >64 | - |
| *Escherichia coli* MO287 | MDR, NDM-1<sup>+</sup> | >64 | - |
| *Escherichia coli* SI-1896 | MDR, OXA-48-like<sup>+</sup> | >64 | - |
| *Klebsiella pneumoniae* B2 | pandrug-resistant | >64 | - |
| *Klebsiella pneumoniae* BO4 | MDR, colistin<sup>R</sup> | >64 | - |
| *Klebsiella pneumoniae* SI-518 | MDR, NDM-1<sup>+</sup> | >64 | - |
| *Klebsiella pneumoniae* SI-703A | MDR, OXA-48-like<sup>+</sup> | >64 | - |
| *Pseudomonas aeruginosa* SI-270 | MDR, colistin<sup>R</sup> | >64 | - |

## 4. Conclusions

Herein we report the discovery and structural elucidation of lunaemycins, new cyclic hexapeptide antibiotics only produced by *S. lunaelactis* species that possess the linear plasmid pSLUN1a. The potent antibacterial activity of lunaemycins against antibiotic resistant Gram-positive bacteria may in part explain how *S. lunaelactis* strains have found their own place in extremely oligotrophic and competitive environments such as moonmilk deposits. Next to the proposed structure of 15 lunaemycin derivatives, we also propose the biosynthetic pathway for lunaemycin A production, the main compound of the *lun* BGC of *S. lunaelactis* MM109^T^ (**Figure 9**).

**Figure 9.**
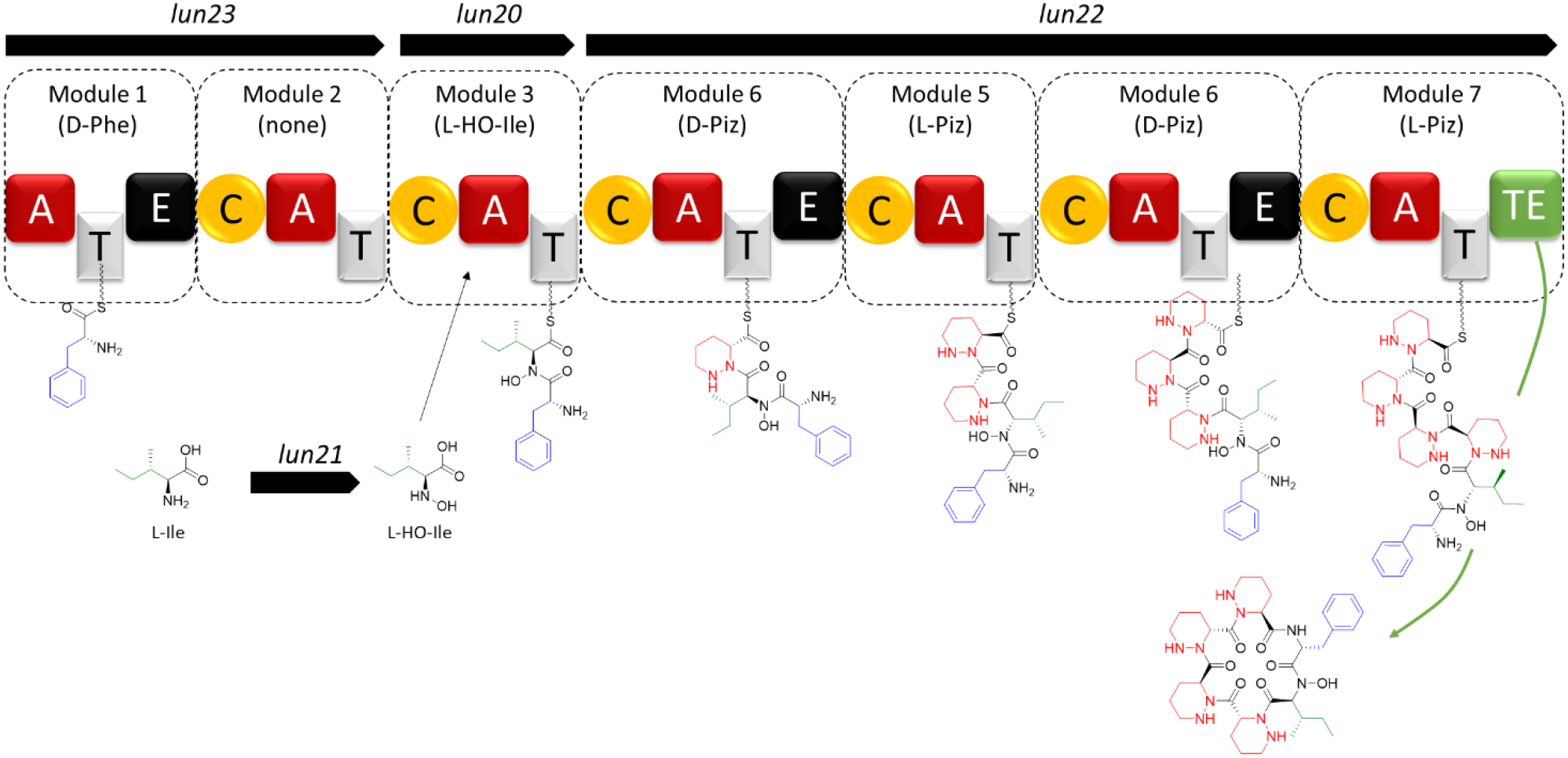
Proposed biosynthetic pathway of lunaemycin A. This model suggests that the Ile will be either N-hydroxylated before activation and recruitment by the A domain (like proposed for the coelichelin pathway) or hydroxylated once activated and tethered on the thiolation domain of Lun20 (like proposed for polyoxypeptin synthesis).

Our work provides a good example where deep *in silico* analysis of genes that compose a BGC combined with exhaustive literature overview of homologous genes allowed to maximize the accuracy of the predicted structure of a cryptic compound. This *in silico* approach highly facilitated the comparative mass spectrometry analysis by guiding the examination on a limited series of ions that possessed the appropriate fragmentation tags. Next to core biosynthetic genes, the *lun* BGC provides also an interesting case where genes required for the synthesis of the building block piperazic acid from ornithine are also present within the BGC itself, and even also the genes that convert the glutamate and glutamine precursors to ornithine. Such situation was previously observed, notably in the BGC for quinichelin biosynthesis **[52]**, and in many NRPS BGCs that necessitate Piz as building block [37]. This inclusion of genes/enzymes of metabolic pathways of primary metabolism within a BGC is most likely mandatory to provide sufficient building blocks at the proper timing of specialized metabolite production.

Another main take-home message of our work relates to the use of multiple strains of a same species to facilitate the correlation between the genetic material associated with the biosynthesis of natural compounds. The current tendency in strain prioritization is instead to try to exclude from a collection the multiple strains that belong to the same species in order to avoid the identification of the same bioactive molecules **[22]**. Here we showed how strain redundancy instead facilitated our work, even avoiding the necessity to generate null mutants to link a BGC to its natural product. This is particularly interesting when a BGC is present on a linear plasmid whose presence in multiple copies complicates gene inactivation (about three copies for pSLUN1a in *S. lunaelactis* MM109^T^ **[21]**). Our attempts to delete genes of BGC 28a have been unsuccessful so far. We are currently applying the same strategy to identify and assess the biological activity of the cryptic products of BGC 28b found in the linear plasmid pSLUN1b only present in the second subgroup of *S. lunaelactis* strains **[22]**.

## 4. Materials and Methods

### 4.1. Bacterial strains and culture conditions

All bacterial strains used in this study are listed in **Table 5**. The archetype strain *Streptomyces lunaelactis* MM109^T^ [20,21] and all other *S. lunaelactis* strains used in this study were isolated from moonmilk deposits of the cave ‘la grotte des collemboles” (Comblain-au-Pont, Belgium) [9,10]. *Streptomyces* spores and mycelium stocks were prepared as described in **[53]**. Cultivation media were prepared as described in **[53]** and inoculated plates were incubated for a week at 28°C.

**Table 5.**
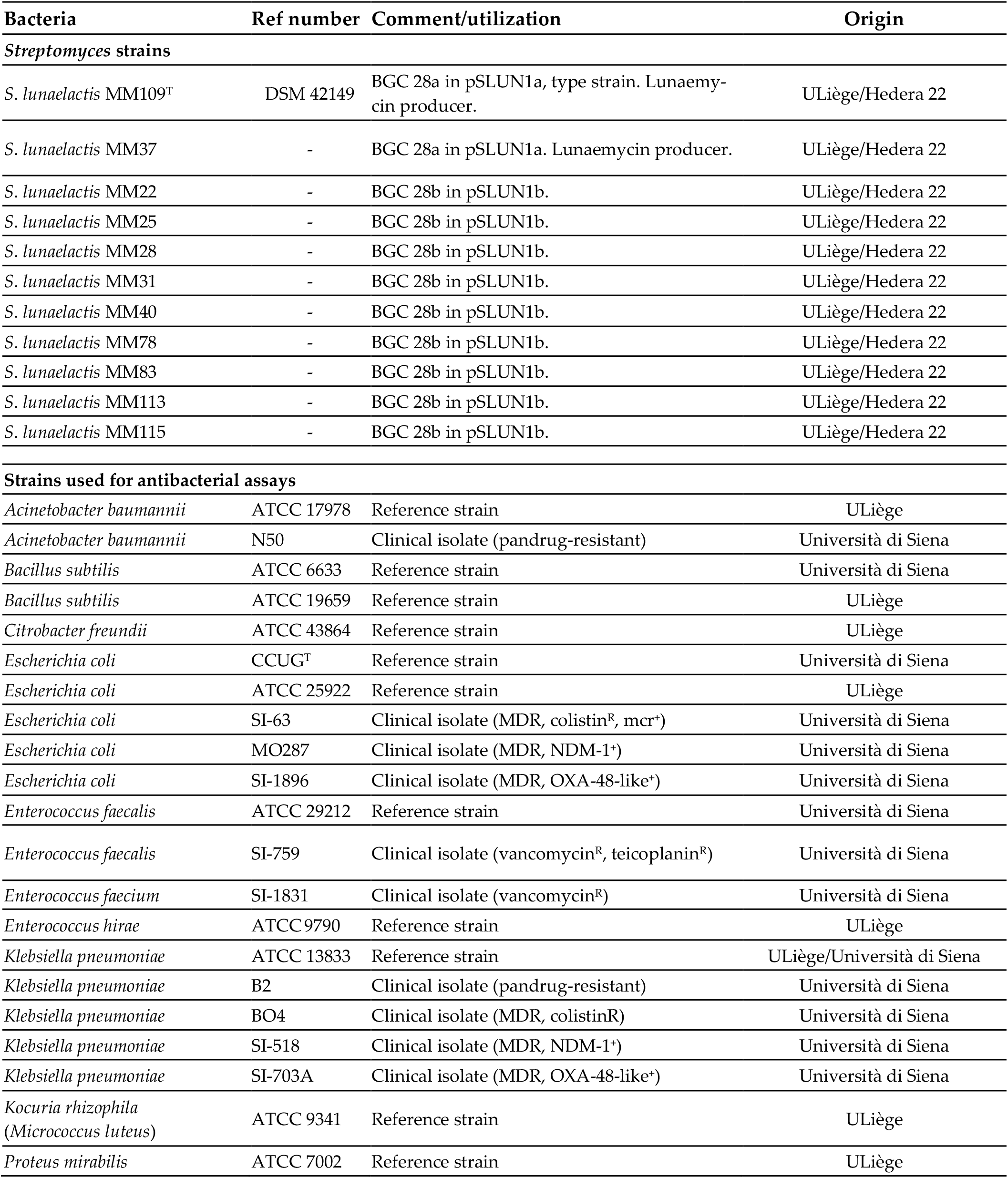
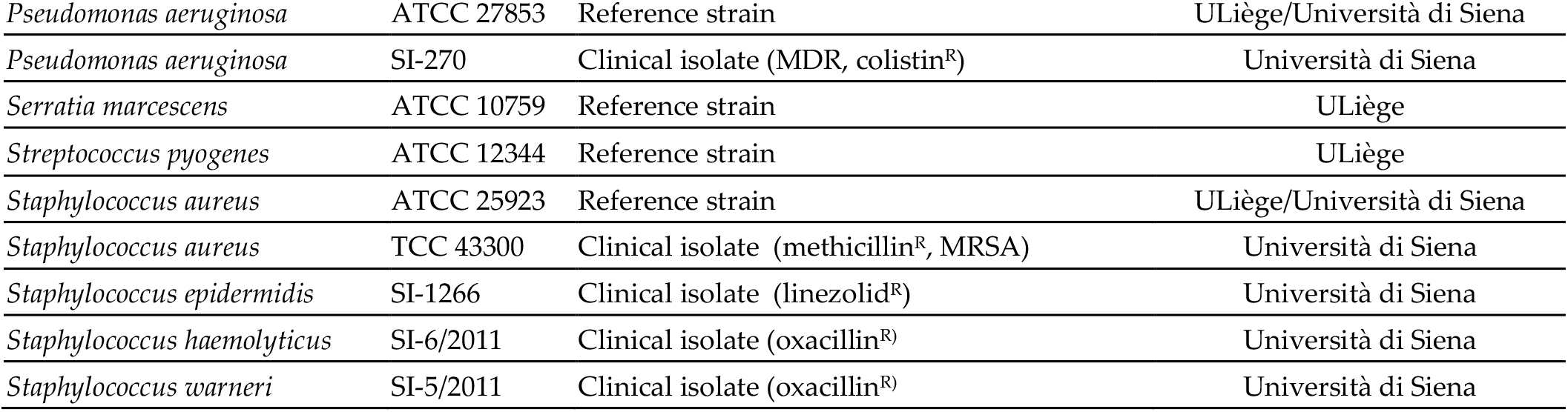
List of bacterial strains used in this study.

### 4.2. Bioinformatics and *in silico* analyzes

Mining for BGC in pSLUN1a (NZ_CP026305.1) and identification of closest homologous genes in known BGCs in MiBIG database **[27]** were performed with antiSMASH **[54]**. Sequence of linear plasmid pSLUN1b can be accessed from (https://www.ncbi.nlm.nih.gov/search/all/?term=JABSUF010000009). Prediction of amino acid building blocks recruited by adenylation domains was performed with SANDPUMA **[39]** and NRPSpredictor2 **[55]**. Visualization of the molecular network (metabolite diversity and relatedness)) was performed via the Global Natural Products Social Molecular Networking website (GNPS, https://gnps.ucsd.edu/) from the tandem mass (MS/MS) spectrometry data obtained from the culture extracts od the different S. *lunaelactis* strains as previously described **[22]**. 1D- and 2D NMR data were analyzed with MestreNova V.14 (https://mestrelab.com/download_file/mnova-14-0-0/).

### 4.3. Compound identification by Ultra-Performance Liquid Chromatography–Tandem Mass Spectrometry (UPLC–MS/MS)

Extracts were analyzed by Ultra-Performance Liquid Chromatography-High Resolution Mass Spectrometry (UPLC–HRMS) and by Ultra–Performance Liquid Chromatography-Tandem Mass Spectrometry (UPLC–MS/MS). Lunaemycins were identified according to their exact mass, the isotopic pattern, their MS/MS spectrum of the molecular ion HCD fragmentation, and the UV-VIS absorbance spectra. The detailed protocols for lunaemycins extraction and purification are detailed in point 4.4. and appendices A-E.

### 4.4. Nuclear magnetic resonance (NMR)

The production and purification of lunaemycins was performed as follows: the ethyl acetate extract of *S. lunaelactis* MM109^T^ grown on 25 agar plates of ISP1 + GlcNAc (~500 ml volume) was dried with rotary evaporator (Rotavapor^®^ R-100) which allowed to obtain 126 mg of a pale yellow powder, re-dissolved in 60 ml of an acetonitrile/water (MilliQ filtered) (10:3, v/v) solution, and fractioned on a C18 LC column (Phenomenex) using an ÄKTA™ purification system. Metabolite elution was performed by using a gradient of increasing concentration of acetonitrile (LC method, appendix A), and each of the 90 fractions were analyzed by HPLC (HPLC method, appendix B) to detect peaks corresponding to lunaemycins. Fractions containing the compounds of interest (fractions 35, 36, and 37 to 41) were subjected to a second round of purification using semi-preparative HPLC (SP-HPLC method, appendix C). Fractions 37 to 41 were pooled, and the volume of each fraction was reduced to 2 ml by evaporation (under gas flow) which served for twenty semi-preparative HPLC injections with 100 μl of each fraction. From fraction 35 we generated 21 sub-fractions among which 4 of them contain lunaemycin D (Table 3) as subsequent UPLC-HRMS analysis (UPLC Method appendix D, and the MS Method, appendix E) identified the molecular ion species of *m/z* 739.39 [M+H]^+^. Fraction 36 generated 19 sub-fractions among which sub-fractions 3 to 5 contain lunaemycin D (previously identified in fraction 35), and sub-fractions 17 to 19 contain lunaemycin B1 as confirmed by UPLC-HRMS analysis that detected the molecular ion species of *m/z* 723.40 [M+H]^+^. Finally, the pooled fractions 37-41 generated 40 sub-fractions (SP-HPLC method, appendix C), among which sub-fractions 23 to 28 mainly contain lunaemycin A (ion species of *m/z* 725.41 [M+H]^+^, see Figure 4a), and lower amounts of lunaemycins B1/B2 (ion species of *m/z* 723.40 [M+H]^+^, see Figure 4a). The fractions containing a single chromatographic peak were pooled and evaporated under continuous gas flow and resuspended in 1 ml of water/acetonitrile (1/9, v/v) solution and diluted to reach a concentration of 0.1 mg/ml for subsequent antibacterial assays (see point 4.5 below).

The pure dried extract mainly containing lunaemycins A (compound **1**) was dissolved in 250 μl of hexadeutero-dimethylsulfoxide (DMSO-d6) containing 1% (v/v) of tetramethylsilane (TMS) as a reference (Sigma-Aldrich), and transferred in a 3 mm NMR tube (103.5 mm long, Bruker). NMR spectra were acquired using a Bruker Avance NEO 700 MHz spectrometer equipped with a cryoprobe and Bruker’s TopSpin 3.5 software package, in order to perform ^1^H, ^13^C (APT), COSY, ^1^H-^13^C HMBC and ^1^H-^13^C HSQC experiments (standard Bruker parameters) and data were interpretated using the MestreNova V.14 software. Sample temperatures were controlled with the variable-temperature unit of the instrument.

### 4.5. Antibacterial assays

Pure lunaemycins A, B1, and D were obtained as described above in point 4.4. Agar-diffusion tests were implemented as described previously **[24]**. Briefly, few microliters of an overnight culture (10 μl of the bacterial stock spread on the solid LB or BH media) are used to make bacterial suspensions with standardized turbidity (OD600 between 0.08 and 0.1, corresponding to a 0.5 McFarland turbidity standard). 100 μl of the standardized bacterial suspension are then inoculated on solid LB or BH media, and 3 mm thick Whatman^®^ papers discs (Sigma-Aldrich, St. Louis, MO, USA) containing 10 μl (2 × 5 μl) of antibiotics are placed on the freshly inoculated solid medium. After overnight incubation at 37°C, the antibacterial activity is evaluated by measuring the diameter of growth inhibition. Control conditions were performed using gentamycin 100 μg/ml as a positive control (corresponding to 1 μg of gentamycin placed on the positive control disc), and acetonitrile, DMSO or water as a negative control. Additional antibacterial susceptiblity assays were performed at the High-Throughput Screening and Protein Engineering platform (HiProtEn, Dipartimento di Biotecnologie Mediche – Università di Siena). Minimal inhibitory concentration (MIC) and minimal bactericidal concentration (MBC) values were determined in Mueller-Hinton broth as recommended by CLSI (document M07-A10, 2015 **[56]**), and the European Committee on Antimicrobial Susceptibility Testing (EUCAST document “Terminology relating to methods for the determination of susceptibility of bacteria to antimicrobial agents”, 2000 **[57]**) on reference (type) strains and clinical isolates as indicated (**Table 5**).

## Supplementary Materials

The following supporting information can be downloaded at: www.mdpi.com/xxx/s1, **Figure S1:** Synteny and gene conservation between BGC 28a of *S. lunaelactis* MM109^T^ and its closest known BGCs.; **Figure S2**: ^1^H NMR spectrum of lunaemycin A in DMSO-*d6*.; **Figure S3:** ^1^H-^1^H COSY NMR spectrum of lunaemycin A in DMSO-*d6*.; **Figure S4:** ^1^H-^13^C HSQC NMR spectrum of lunaemycin A in DMSO-*d6*.; **Figure S5:** ^1^H-^13^C HMBC NMR spectrum of lunaemycin A in DMSO-*d6*.; **Figure S6:** ^13^C APT NMR spectrum of lunaemycin A in DMSO-*d6*. Quaternary C/CH2 in negative and CH/CH3 in positive.; **Figure S7:** HRMS/MS spectrum of lunaemycin B1 (compound **2a**, see Table 3 and Figure 7).; **Figure S8:** HRMS/MS spectrum of lunaemycin B2 (compound **2b**, see Table 3 and Figure 7).; **Figure S9:** HRMS/MS spectrum of lunaemycin C1 (compound **3a**, see Table 3 and Figure 7).; **Figure S10:** HRMS/MS spectrum of lunaemycin C2 (compound **3b**, see Table 3 and Figure 7).; **Figure S11:** HRMS/MS spectrum of lunaemycin D (compound **4**, see Table 3 and Figure 7).; **Figure S12:** HRMS/MS spectrum of lunaemycin E1 (compound **5a**, see Table 3 and Figure 7).; **Figure S13:** HRMS/MS spectrum of lunaemycin E2 (compound **5b**, see Table 3 and Figure 7).; **Figure S14:** HRMS/MS spectrum of lunaemycin F (compound **6**, see Table 3 and Figure 7).; **Figure S15:** HRMS/MS spectrum of lunaemycin G (compound **7**, see Table 3 and Figure 7).; **Figure S16:** HRMS/MS spectrum of lunaemycin H (compound **8**, see Table 3 and Figure 7).; **Figure S17:** HRMS/MS spectrum of lunaemycin I (compound **9**, see Table 3 and Figure 7).; **Figure S18:** HRMS/MS spectrum of lunaemycin J1 (compound **10a**, see Table 3 and Figure 7).; **Figure S19:** HRMS/MS spectrum of lunaemycin J2 (compound **10b**, see Table 3 and Figure 7).; **Figure S20:** HRMS/MS spectrum of lunaemycin J3 (compound **10c**, see Table 3 and Figure 7).

## Author Contributions

Conceptualization, L.M. and S.R.; methodology, L.M.; software, A.N., L.R., G.M., and M.F.; validation, L. M., A.N., L.R., B.M., M.F., and S.R.; formal analysis, L.M., A.N.; investigation, L.M., A.N., L.R., D.T., F.S., D.B.; resources, G.M., M. F., and S.R.; Data curation, L.M., A.N, L.R., G.M., M.F., and S.R.; writing — original draft preparation, L.M. and S.R.; writing-review and editing, L.M, L.R., D.T., M.F., D.B., JD.D., and S.R.; visualization, L.M., A.N., L.R., and S.R.; supervision, S.R.; project administration, S.R.; funding acquisition, S.R. All authors have read and agreed to the published version of the manuscript.

## Data Availability Statement

The Molecular Networking job in GNPS can be found at https://gnps.ucsd.edu/ProteoSAFe/status.jsp?task=84ea45db68814c94a3b2bad9c26dd807.

## Acknowledgments

We are grateful to D. Adam, B. Deflandre, M. Maciejewska, E. Tenconi, S. Anderssen, and N. Stulanovic at the Streptomyces Genetics and Development group, and everyone at HEDERA-22 for fruitful discussions and technical support.

## Appendix

### Appendix A - LC method

After equilibration of the stationary phase in 100 ml of a 30% acetonitrile solution, the full extract (diluted in 60 ml of a 30% acetonitrile solution) was loaded on the column, followed by five successive elution steps with 60 ml of a 40, 50, 66, 80, and finally 100% of an acetonitrile solution. In total, 90 fractions were collected (fractions of 5 ml from the full extract injection until the end of the 50% acetonitrile elution step and in the last elution step at 100% of acetonitrile; fractions of 3 ml fractions were collected during the elution steps with 66 and 80% of acetonitrile solution). This method was applied on a GE AKTA Prime Plus Liquid Chromatography System with a bulk of Sepra™ C18-E column (Phenomenex, 50 μm, 65 Å) manually packed in a XK16 bulk (Pharmacia Biotech).

### Appendix B – HPLC method

The HPLC system (HPLC Alliance, Waters) used a reversed phase Luna^®^ Omega PS C18 column (Phenomenex, 2.1mm by 150 mm, 5 μm particle size, 100-Å pore size) column. Metabolites separation was performed using a step gradient system of 0.1% FA in 0.22 μm filtrated milliQ H_2_O (solvent A) and acetonitrile (Barker; HPLC Far UV grade) (solvent B), as follows: after a 2 min equilibration, the gradient started from 5% and went to 100% of B in 80 min, stayed at 100% of B during 8 min, decreased to 5% of B in 8 min and finally pre-equilibrated the system for 2 min; flow rate: 0,5 ml/min; column temperature of 40°C. 50 μl of pre-purified fractions were injected and single UV-Vis absorbance were measured at 195 and 220 nm. Full UV-Vis absorbance scans were also recorded for all analyses (from 190 to 800 nm).

### Appendix C – SP-HPLC method

SP-HPLC column Luna^®^ Omega PS C18 column (Phenomenex, 21.2 mm by 50 mm, 5 μm particle size, 100-Å pore size) allowed accurate purification of lunaemycins using agradient system of 0.1% FA in 0.22 μm filtrated milliQ H_2_O (solvent A) and acetonitrile (Barker; HPLC Far UV grade) (solvent B), as follows: after a 2 min equilibration, the gradient started from 50% and went to 70% of B in 18 min, stayed at 70% of B during 8 min, decreased to 50% of B in 1 min and finally pre-equilibrated the system for 1 min; flow rate: 3.5 ml/min. 50 μl of pre-purified fractions were injected and single UV-Vis absorbance were measured at 195 and 220 nm. Full UV-Vis absorbance scans were also recorded for all analyses (from 190 to 800 nm).

### Appendix D - UPLC method

RP-HPLC column Luna^®^ Omega PS C18 column (Phenomenex, 2.1 mm by 150 mm, 5 μm particle size, 100-Å pore size) allowed the separation of metabolites using a gradient system of 0.1% FA in 0.22 μm filtrated milliQ H_2_O (solvent A) and acetonitrile (Barker; HPLC Far UV grade) (solvent B), as follows: after 2 min of equilibration of the column, the gradient started from 5% and went to 100% of B in 23 min, stayed at 100% of B during 2 min, decreased to 5% of B in 1 min and finally equilibrated the system for 2 min; flow rate: 0.4 ml/min; column temperature 40°C. 10 μl of each fraction were injected and absorbance was measured at 220 and 280 nm.

### Appendix E - MS Method

All HRMS and HRMS/MS spectra were obtained using the same UPLC-HRMS/MS method. High-Resolution Electrospray Ionization Mass Spectrometry - HRESIMS data were acquired on a Q Exactive Plus hybrid Quadrupole-Orbitrap mass spectrometer equipped with an electrospray ionization (ESI) source (Q Exactive Plus, Thermo Fisher Scientific, San Jose, CA, USA). Briefly, compounds were separated by reverse-phase chromatography using ultraperformance liquid chromatography (UPLC I-Class, Waters) as described in Appendix D - UPLC method. This chromatographic system was coupled to a Q Exactive Plus, operated in electrospray positive ion mode and programmed for data-dependent acquisitions. Survey scans were acquired at a mass-resolving power level of 140,000 FWHM (full width at half-maximum) from 100 to 1,500 *m/z* (accumulation target, 1 x 10^6^ ions or maximum inject time of 200 ms). The five ions showing the greatest intensity were then selected for tandem mass spectrometry (MS/MS) experiments by the use of higher-energy collision dissociation (HCD) fragmentation and stepped normalized collision energy (NCE) (21.2; 25; 28) within 2-atomic-mass-unit (amu) isolation window (resolution, 17,500; accumulation target, 1 x 10^5^ ions or maximum inject tim of 200 ms). Dynamic exclusion was enabled for 10 s. Data were analyzed using Xcalibur v2.2 (Thermo Fisher Scientific, San Jose, CA, USA). Each of the compounds was identified in accordance with its exact mass and isotope pattern, observation of the MS/MS spectra obtained by molecular ion fragmentation, and the UV-VIS absorbance.

