## Supplementary Figures for "Lunaemycins, new cyclic hexapeptide antibiotics from the cave moonmilk-dweller *Streptomyces lunaelactis* MM109^T^"

### Supplementary Files.

#### Figure S1. Comparative analysis of BGC 28a and its closest known BGCs.

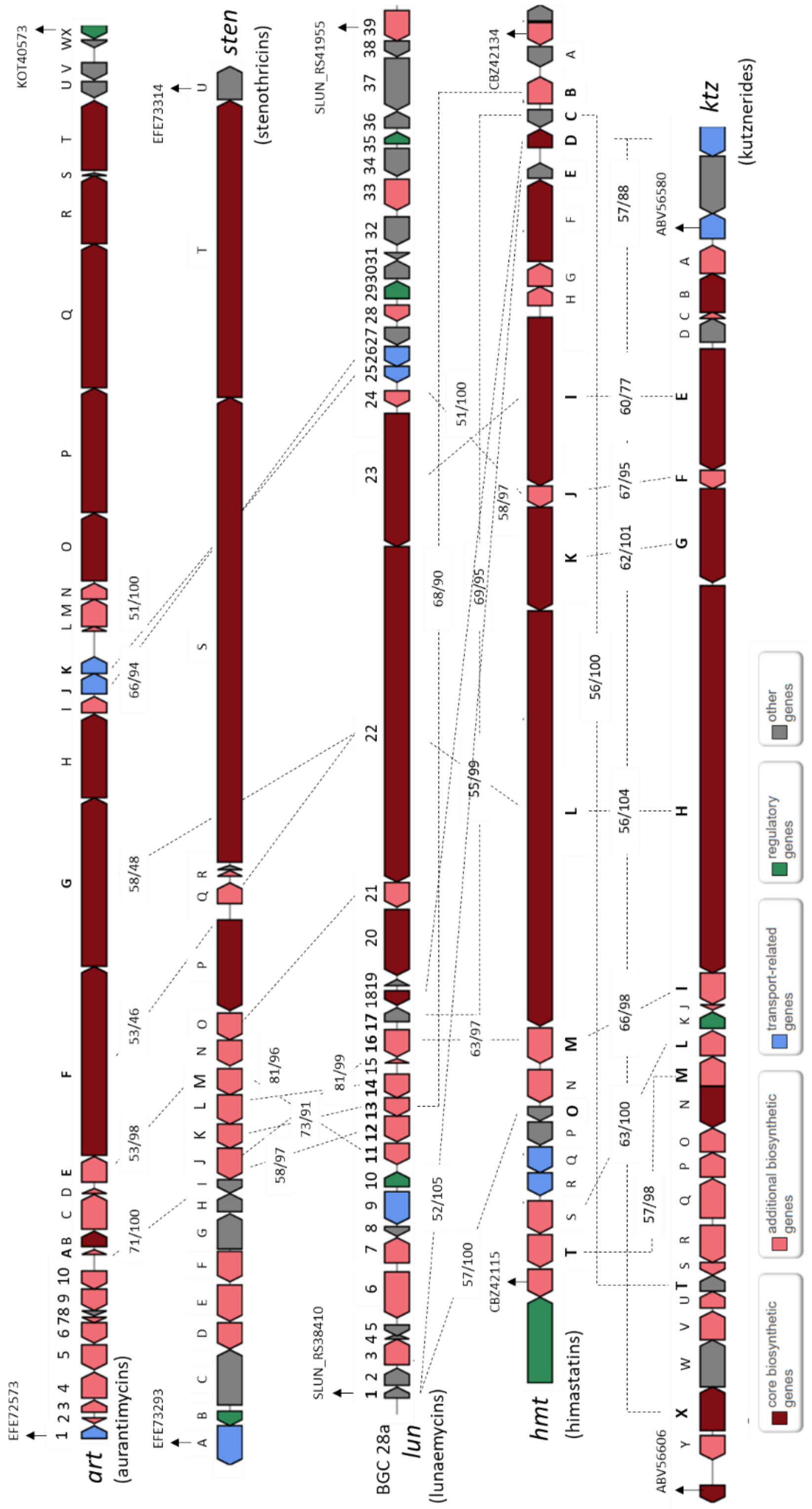


**Figure S1.** Synteny and gene conservation between BGC 28a of *S. lunaelactis* MM109$^{T}$ and its closest known BGCs. Dotted lines link homologous genes between clusters. Numbers associated with each pair of homologous ORFs refer to the percentage of aa identity and the sequence coverage, respectively.

### Figure S2-S6. NMR data.

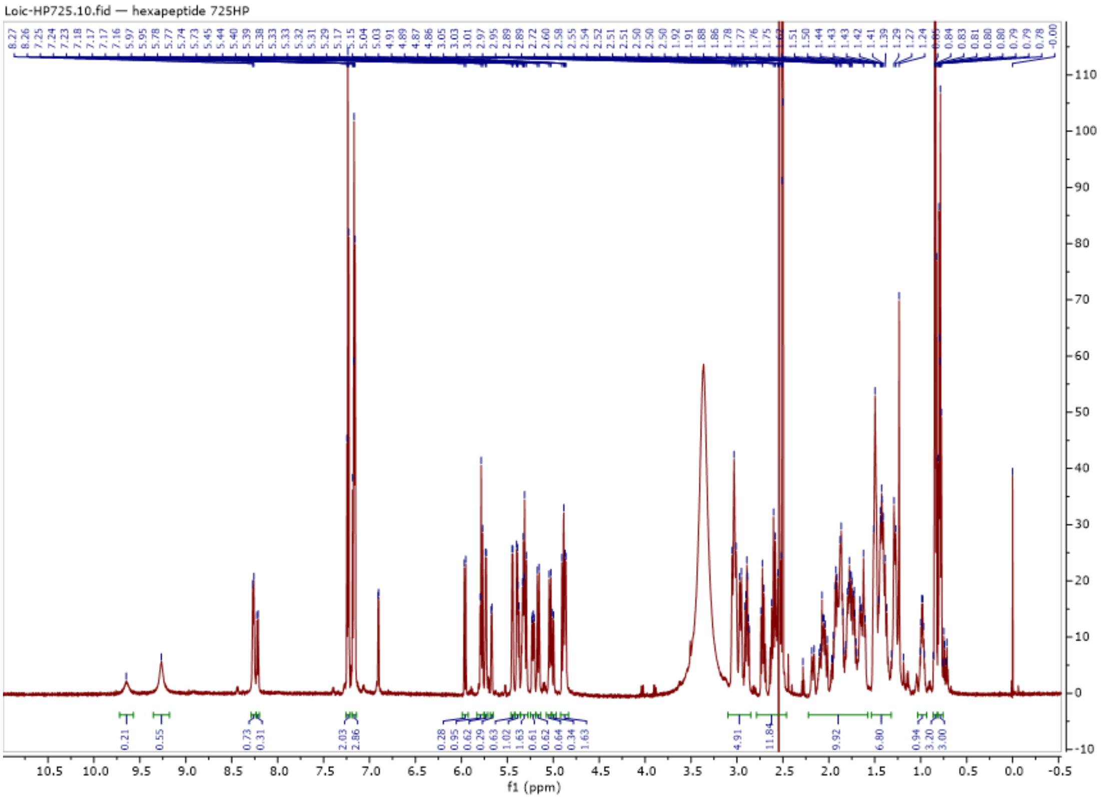


**Figure S2.** $^{1}H$ NMR spectrum of lunaemycin A in DMSO-*d6*.

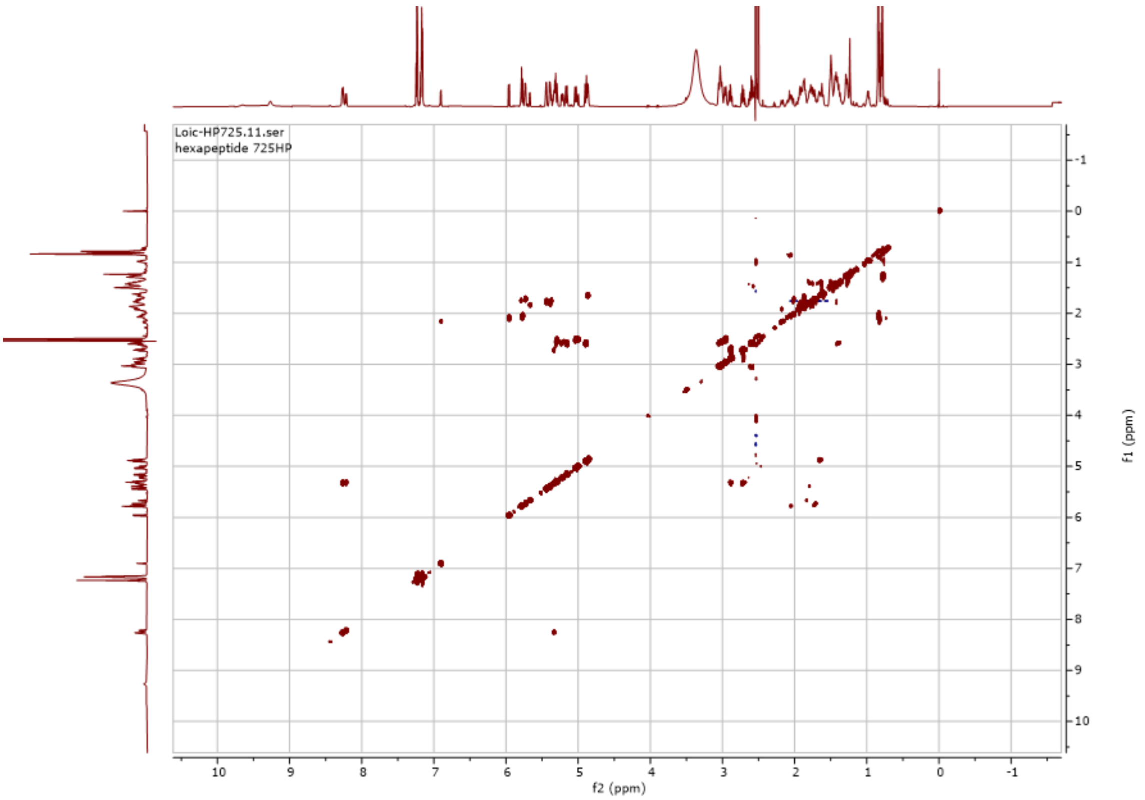


**Figure S3.** $^{1}H$-$^{1}H$ COSY NMR spectrum of lunaemycin A in DMSO-*d6*.

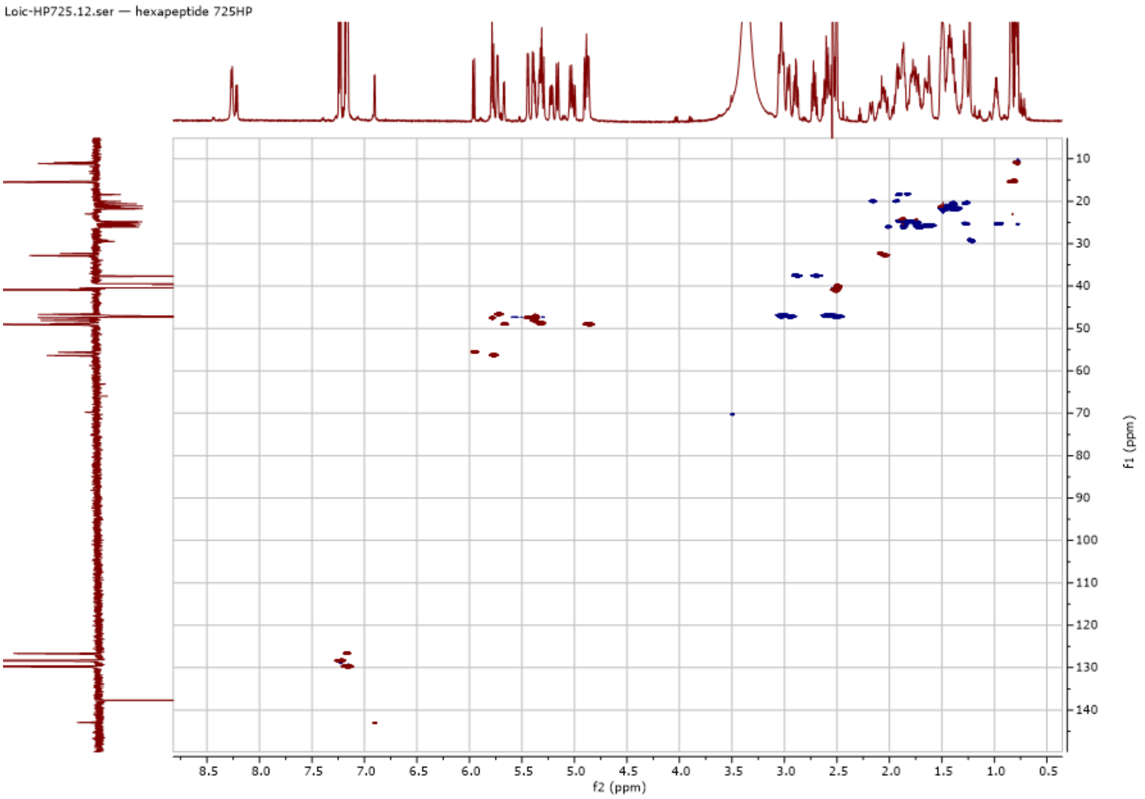


**Figure S4.** $^{1}H$-$^{13}C$ HSQC NMR spectrum of lunaemycin A in DMSO-*d6*.

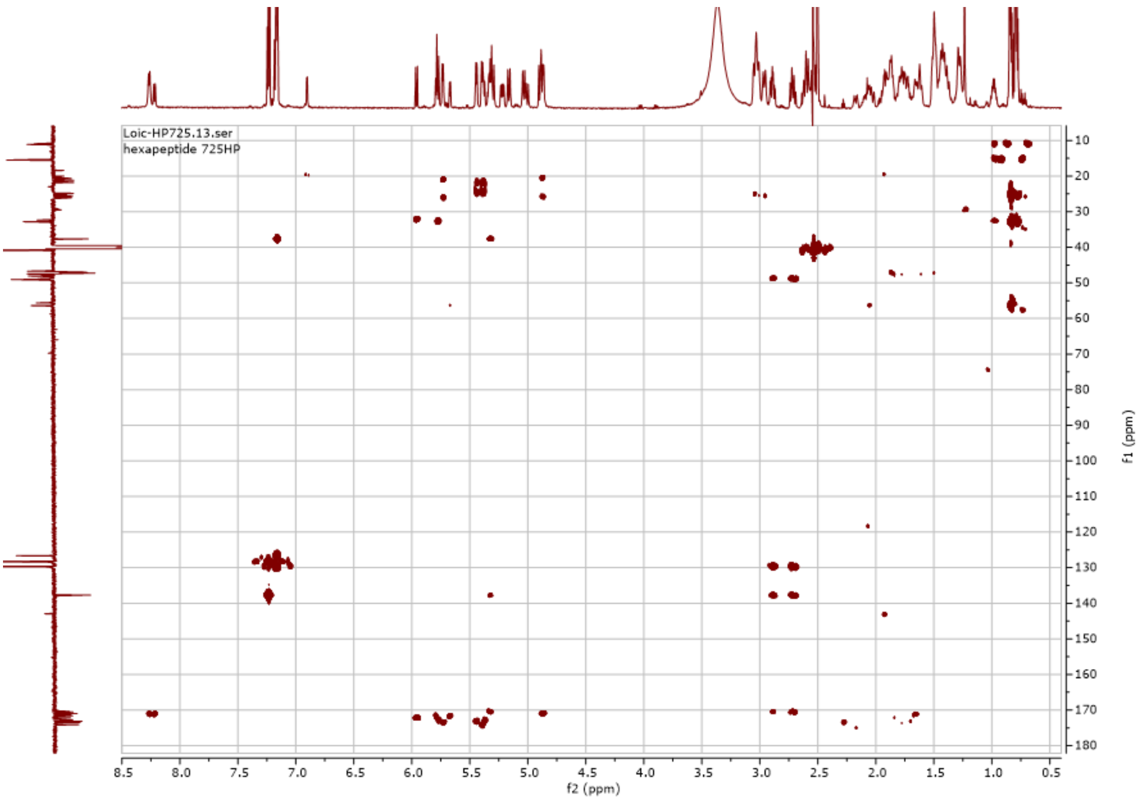


**Figure S5.** $^{1}H$-$^{13}C$ HMBC NMR spectrum of lunaemycin A in DMSO-d6.

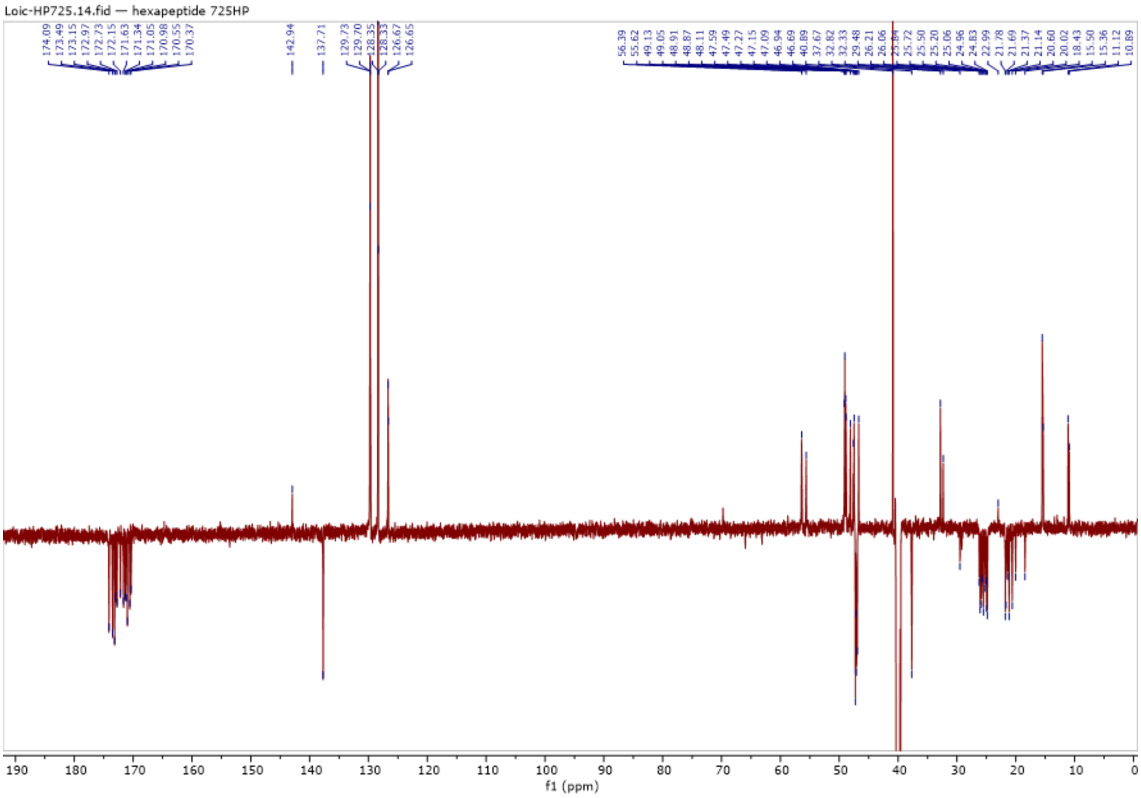


**Figure S6.** $^{13}C$ APT NMR spectrum of lunaemycin A in DMSO-d6. Quaternary C/CH2 in negative and CH/CH3 in positive.

**Figure S7-S20. Proposed structure for the 15 lunaemycins identified in *S. lunaelactis* extracts deduced by HRMS/MS analysis.** Only the fragments corresponding to amino-acids or peptides *m/z* are highlighted on the mass spectrum. Next to each spectrum, the proposed structure and its fragmentation mechanisms are shown. Color code: orange, piperazic acid (Piz); grey, phenylalanine (Phe); light blue, hydroxylated isoleucine (HO-Ile); deep blue, hydroxyl-dehydropiperazate (HOdhPiz); yellow, hydroxyvaline (HO-Val); red corresponds to a fragment containing more than one modified residue compared to lunaemycin A.

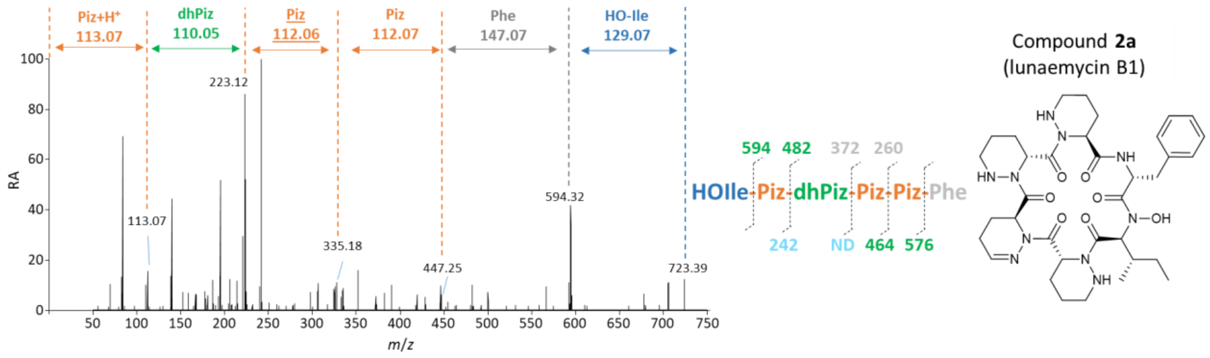


**Figure S7.** HRMS/MS spectrum of lunaemycin B1 (compound **2a**, Table 3, Figure 7).

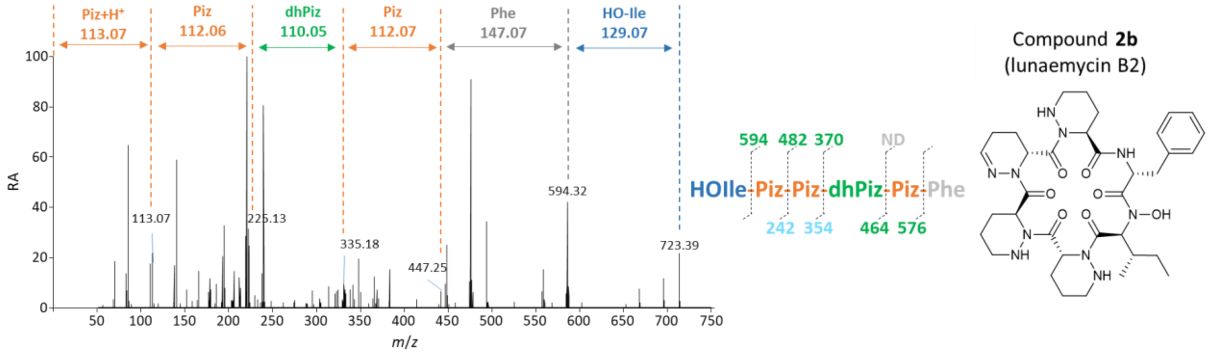


**Figure S8.** HRMS/MS spectrum of lunaemycin B2 (compound **2b**, Table 3, Figure 7).

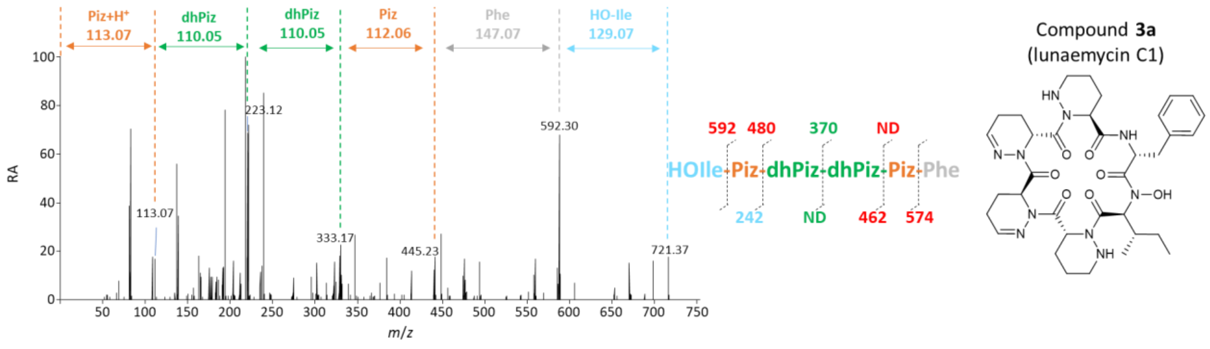


**Figure S9.** HRMS/MS spectrum of lunaemycin C1 (compound **3a**, Table 3, Figure 7).

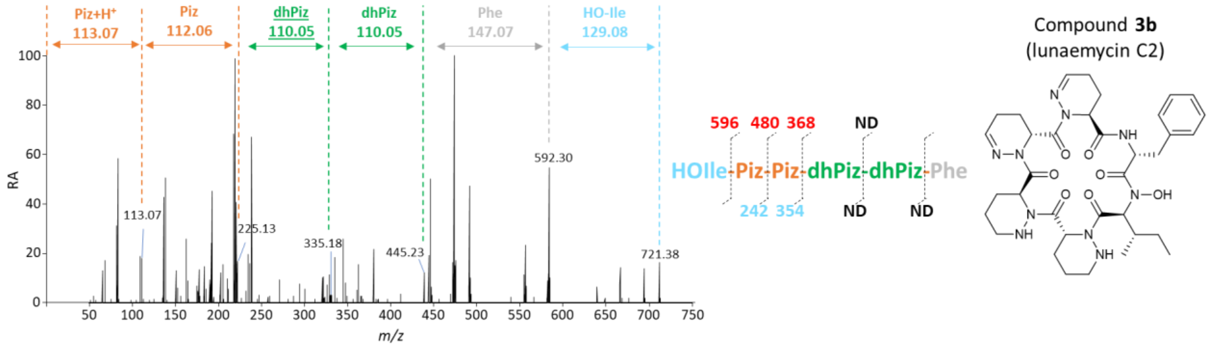


**Figure S10.** HRMS/MS spectrum of lunaemycin C2 (compound **3b**, Table 3, Figure 7).

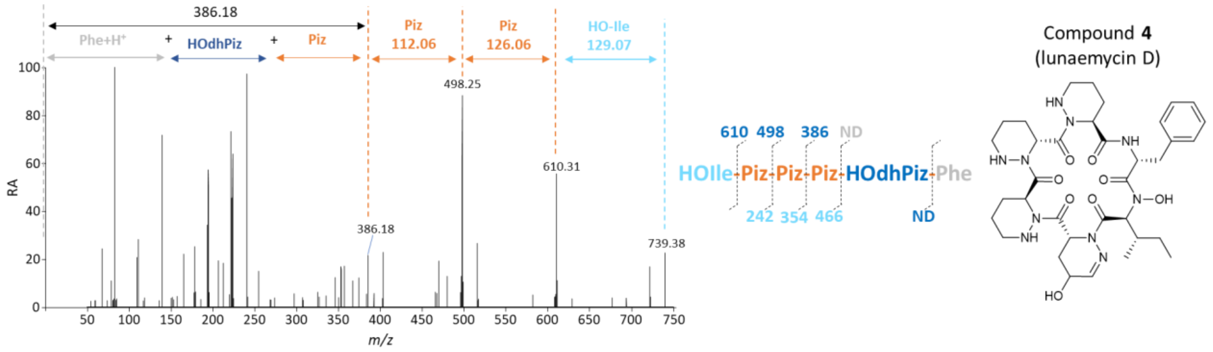


**Figure S11.** HRMS/MS spectrum of lunaemycin D (compound **4**, Table 3, Figure 7).

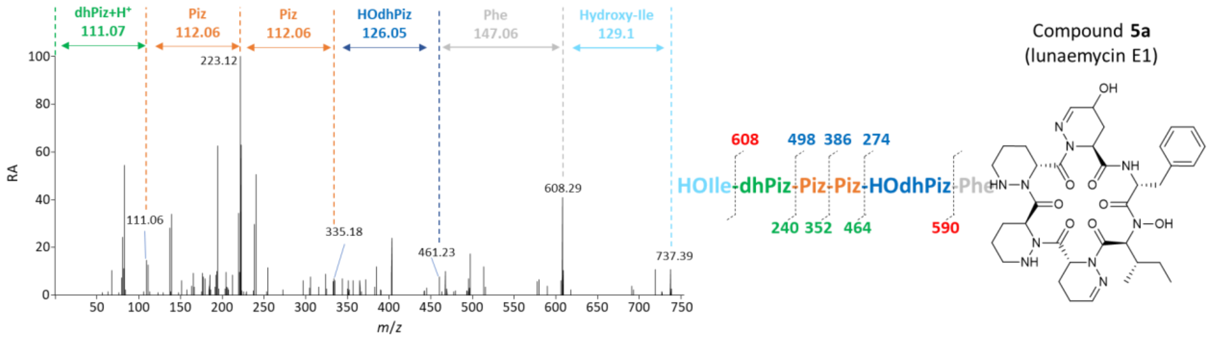


**Figure S12.** HRMS/MS spectrum of lunaemycin E1 (compound **5a**, Table 3, Figure 7).

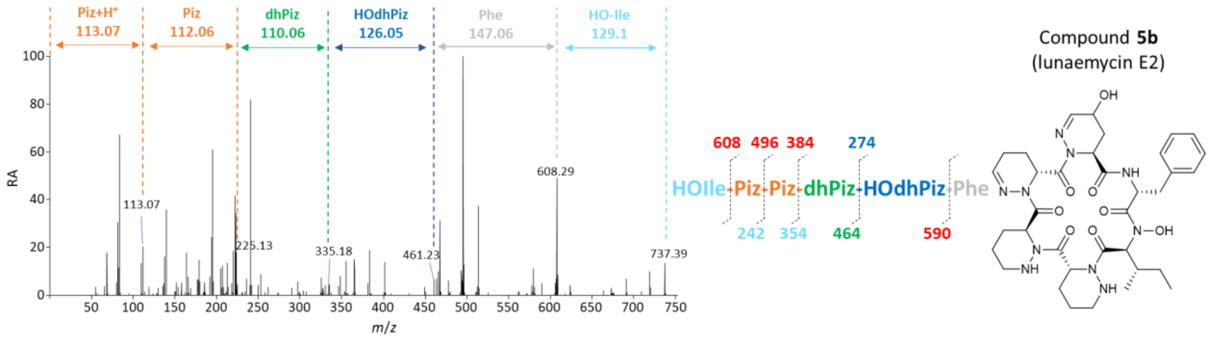


**Figure S13.** HRMS/MS spectrum of lunaemycin E2 (compound **5b**, Table 3, Figure 7).

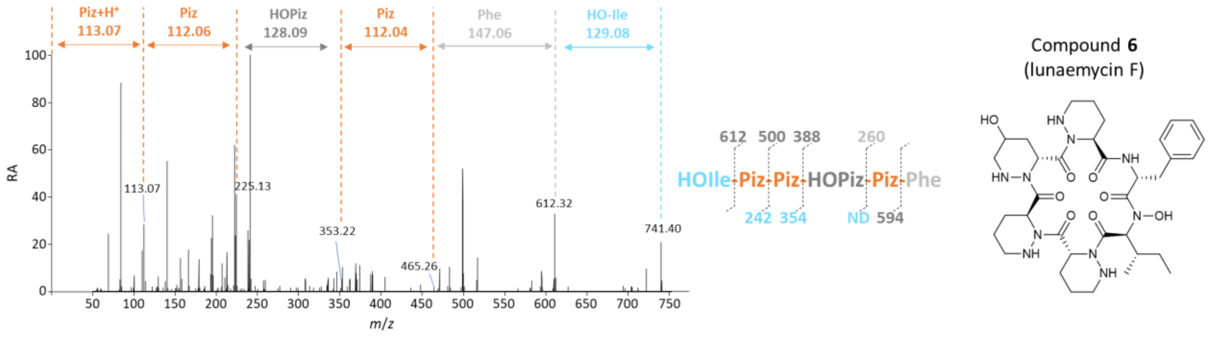


**Figure S14.** HRMS/MS spectrum of lunaemycin F (compound **6**, Table 3, Figure 7).

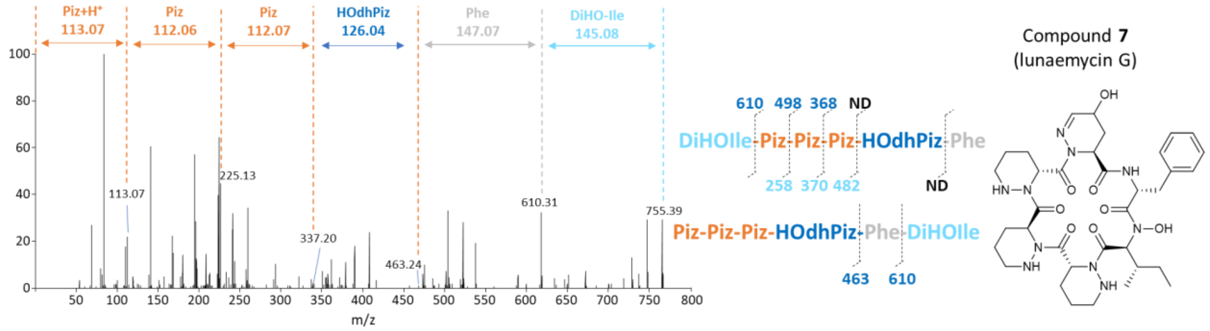


**Figure S15.** HRMS/MS spectrum of lunaemycin G (compound **7**, Table 3, Figure 7).

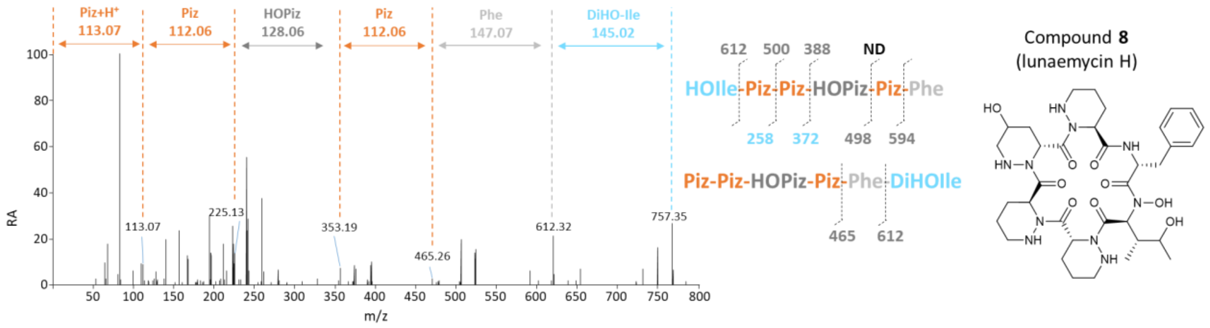


**Figure S16.** HRMS/MS spectrum of lunaemycin H (compound **8**, Table 3, Figure 7).

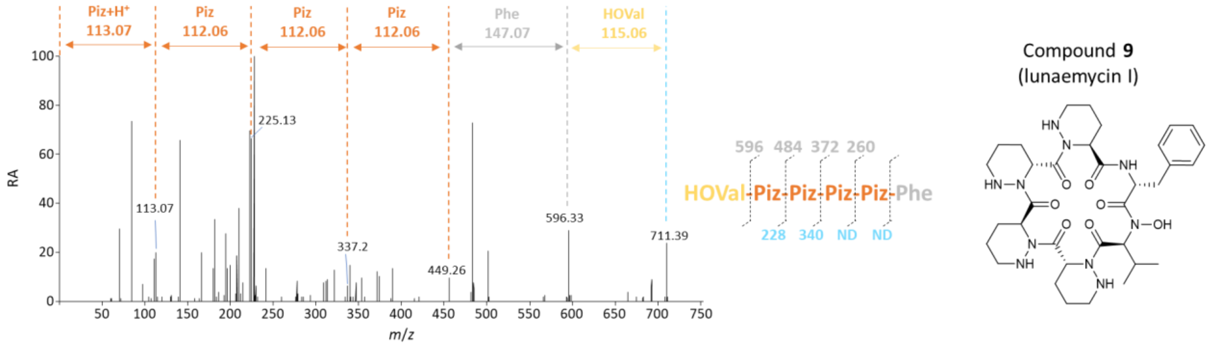


**Figure S17.** HRMS/MS spectrum of lunaemycin I (compound **9**, Table 3, Figure 7).

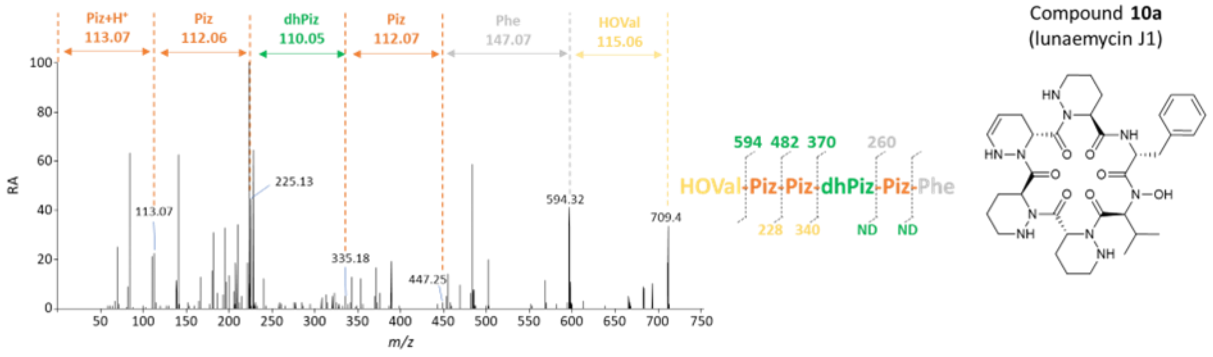


**Figure S18.** HRMS/MS spectrum of lunaemycin J1 (compound **10a**, Table 3, Figure 7).

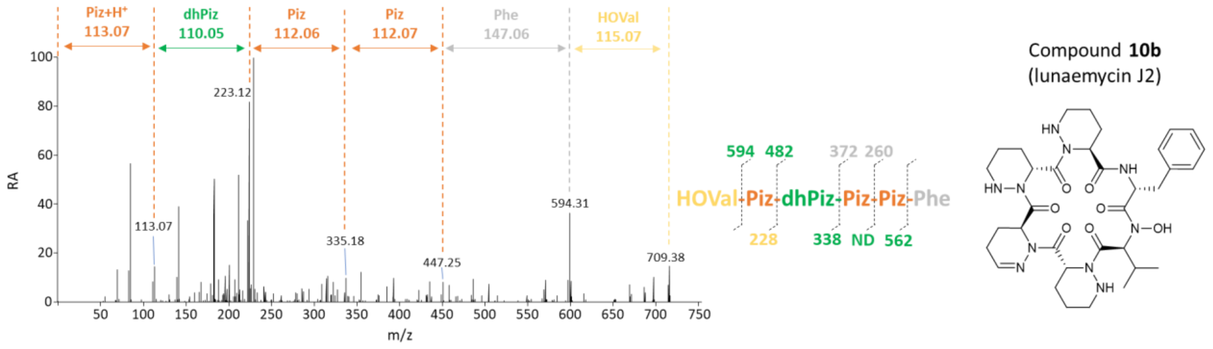


**Figure S19.** HRMS/MS spectrum of lunaemycin J2 (compound **10b**, Table 3, Figure 7).

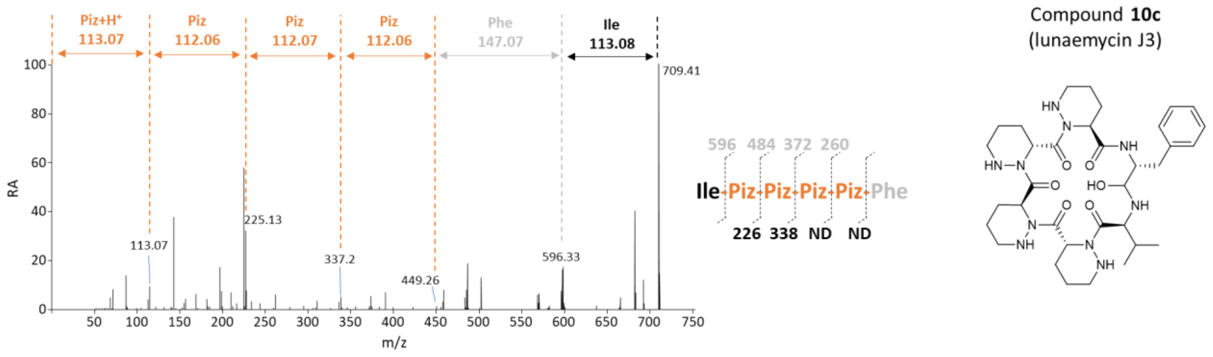


**Figure S20.** HRMS/MS spectrum of lunaemycin J3 (compound **10c**, Table 3, Figure 7).
